# Alpha-arrestins Aly1 and Aly2 regulate trafficking of the glycerophosphoinositol transporter Git1 and impact phospholipid homeostasis

**DOI:** 10.1101/2021.02.04.429748

**Authors:** Benjamin P. Robinson, Sarah Hawbaker, Annette Chiang, Eric M. Jordahl, Sanket Anaokar, Alexiy Nikiforov, Ray W. Bowman, Philip Ziegler, Ceara K. McAtee, Jana Patton-Vogt, Allyson F. O’Donnell

**Affiliations:** Department of Biological Sciences, University of Pittsburgh, Pittsburgh, PA, USA; Department of Biological Sciences, Duquesne University, Pittsburgh, PA, USA

**Author notes:** Corresponding author: Department of Biological Sciences, University of Pittsburgh, 4249 Fifth Avenue, Pittsburgh, PA, USA 15206.

**Keywords:** E3 ubiquitin ligase, endocytosis, *S. cerevisiae*, vacuole, calcineurin, microscopy, protein trafficking adaptors

## Abstract

Phosphatidylinositol (PI) is an essential phospholipid and critical component of membrane bilayers. The complete deacylation of PI by phospholipases of the B-type leads to the production of intracellular and extracellular glycerophosphoinositol (GPI), a water-soluble glycerophosphodiester. Extracellular GPI is transported into the cell via Git1, a member of the Major Facilitator Superfamily of transporters that resides at the plasma membrane in yeast. Once internalized, GPI can be degraded to produce inositol, phosphate and glycerol, thereby contributing to reserves of these building blocks. Not surprisingly, *GIT1* gene expression is controlled by nutrient balance, with limitation for phosphate or inositol each increasing *GIT1* expression to facilitate GPI uptake. Less is known about how Git1 protein levels or localization are controlled. Here we show that the α-arrestins, an important class of protein trafficking adaptor, regulate the localization of Git1 in a manner dependent upon their association with the ubiquitin ligase Rsp5. Specifically, α-arrestin Aly2 is needed for effective Git1 internalization from the plasma membrane under basal conditions. However, in response to GPI-treatment of cells, either Aly1 or Aly2 can promote Git1 trafficking to the vacuole. Retention of Git1 at the cell surface, as occurs in *aly1*Δ *aly2*Δ cells, results in impaired growth in the presences of excess exogenous GPI and results in increased uptake of radiolabeled GPI, suggesting that accumulation of this metabolite or its downstream products leads to cellular toxicity. We further show that regulation of α-arrestin Aly1 by the protein phosphatase calcineurin improves both steady-state and ligand-induced trafficking of Git1 when a mutant allele of Aly1 that mimics the dephosphorylated state at calcineurin-regulated residues is employed. Thus, calcineurin regulation of Aly1 is important for the GPI-ligand induced trafficking of Git1 by this α-arrestin, however, the role of calcineurin in regulating Git1 trafficking is much broader than can simply be explained by regulation of the α-arrestins. Finally, we find that loss of Aly1 and Aly2 leads to an increase in phosphatidylinositol-3-phosphate on the limiting membrane of the vacuole and this alteration is further exacerbated by addition of GPI, suggesting that the effect is at least partially linked to Git1 function. Indeed, loss of Aly1 and Aly2 leads to increased incorporation of inositol label from ^3^H-inositol-labelled GPI into PI, confirming that internalized GPI influences PI synthesis and indicating a role for the α-arrestins in regulating the process.

## Introduction

Maintaining nutrient balance is critical for cellular function. Eukaryotic cells are divided into membrane-bound organelles, each of which has a unique lipid composition that helps define the organelle, alters its function and influences protein trafficking to and from these compartments (Behnia and Munro, 2005; Strahl and Thorner, 2007; Klug and Daum, 2014; Casares *et al*., 2019). The lipid bilayers surrounding organelles contain membrane-embedded transporters that allow for nutrient and metabolite entry. Specific transporters reside in the plasma membrane (PM)—allowing for compounds to enter the cell and controlling the exit of toxic metabolites—and in the membranes surrounding other organelles, allowing the balance of intracellular metabolites to be fine-tuned as required for cellular health. Importantly, imbalances in lipids, nucleotides or amino acids often results in cellular toxicity (Kim *et al*., 2000; Risinger *et al*., 2006; Ljungdahl and Daignan-Fornier, 2012; Shyu *et al*., 2019; Ruiz *et al*., 2020). Here we focus on regulation of the glycerophosphoinositol (GPI) transporter and consider how elevated GPI transport, which leads to increased cellular phosphatidylinositol (PI), may alter downstream pathways (Patton *et al*., 1995; Patton-Vogt and Henry, 1998).

During the course of normal metabolism, PI is deacylated by phospholipases of the B-type to produce extracellular GPI (Angus and Lester, 1972; Patton *et al*., 1995). GPI is a major product of PI turnover, the other being the complex inositol-containing sphingolipids (Angus and Lester, 1972). GPI, and to a lesser extent, glycerophosphocholine, can then enter the cell via the transporter encoded by the *GIT1* gene (Patton-Vogt and Henry, 1998; Fisher *et al*., 2005). Once internalized, GPI is eventually broken down into inositol, phosphate and glycerol, thus contributing to the cellular pool of these metabolites (Patton-Vogt and Henry, 1998). In fact, Git1 was first cloned for its ability to restore growth to the inositol auxotroph, *ino1*, which harbors a mutation in inositol-1-phosphate synthase (Patton-Vogt and Henry, 1998), demonstrating that uptake of GPI through Git1 provides inositol to cells (Almaguer *et al*., 2003; Almaguer *et al*., 2004). Not surprisingly, transcription of the *GIT1* gene is regulated by inositol, and more exquisitely by phosphate (Almaguer *et al*., 2003; Almaguer *et al*., 2004). The *GIT1* promoter contains binding sites for the transcription factors Pho2 and Pho4 (Almaguer *et al*., 2004), which respond to phosphate limitation by co-operatively binding to the promoters of genes encoding proteins needed for phosphate uptake and/or scavenging (Vogel *et al*., 1989; Magbanua *et al*., 1997; Shao *et al*., 1998). The *GIT1* promoter further contains inositol and choline responsive elements (ICREs) for the heteromeric Ino2/Ino4 transcription factors, consistent with a role for inositol restriction in stimulating *GIT1* gene expression. However, inositol limitation plays a more modest role in the amplitude of *GIT1* gene expression than phosphate starvation (Almaguer *et al*., 2004).

Once GPI enters the cell through Git1, the release of free inositol fluxes into PI, which in turn, is a precursor to two important classes of lipids: polyphosphoinositides and sphingolipids (Figure 1A) (Angus and Lester, 1972; Patton *et al*., 1995). Phosphoinositides are an important but minor class of membrane lipids produced by phosphorylation/dephosphorylation of the inositol ring of PI. Specifically, myo-inositol can be phosphorylated on the 3, 4 and 5 carbon positions, and each organelle and/or subdomain is enriched for a specific class of phosphorylated PI (Di Paolo and De Camilli, 2006; Strahl and Thorner, 2007). The lipid kinases and phosphatases involved are in turn controlled by signaling events that direct their activity or localization to allow specific phospholipid species to be made or turned over (Strahl and Thorner, 2007). These phospholipids are important binding sites at the membrane surface that regulate many membrane trafficking and signaling events. In contrast, the sphingolipids are a major lipid species in yeast membranes with key roles in regulating endocytosis, actin organization, response to heat stress, calcium sensitivity and cell cycle regulation (Klemm *et al*., 2009; Epstein *et al*., 2012; Klug and Daum, 2014; Montefusco *et al*., 2014). PI is used in *de novo* sphingolipid synthesis at two steps in the complex sphingolipid base formation (see Figure 8C) (Dickson *et al*., 2006; Klug and Daum, 2014; Montefusco *et al*., 2014); therefore, perturbations in PI can alter sphingolipid synthesis in cells, with increasing PI favoring elevated flux through the sphingolipid biosynthetic pathway (Henry *et al*., 2012).

**FIGURE 1.**
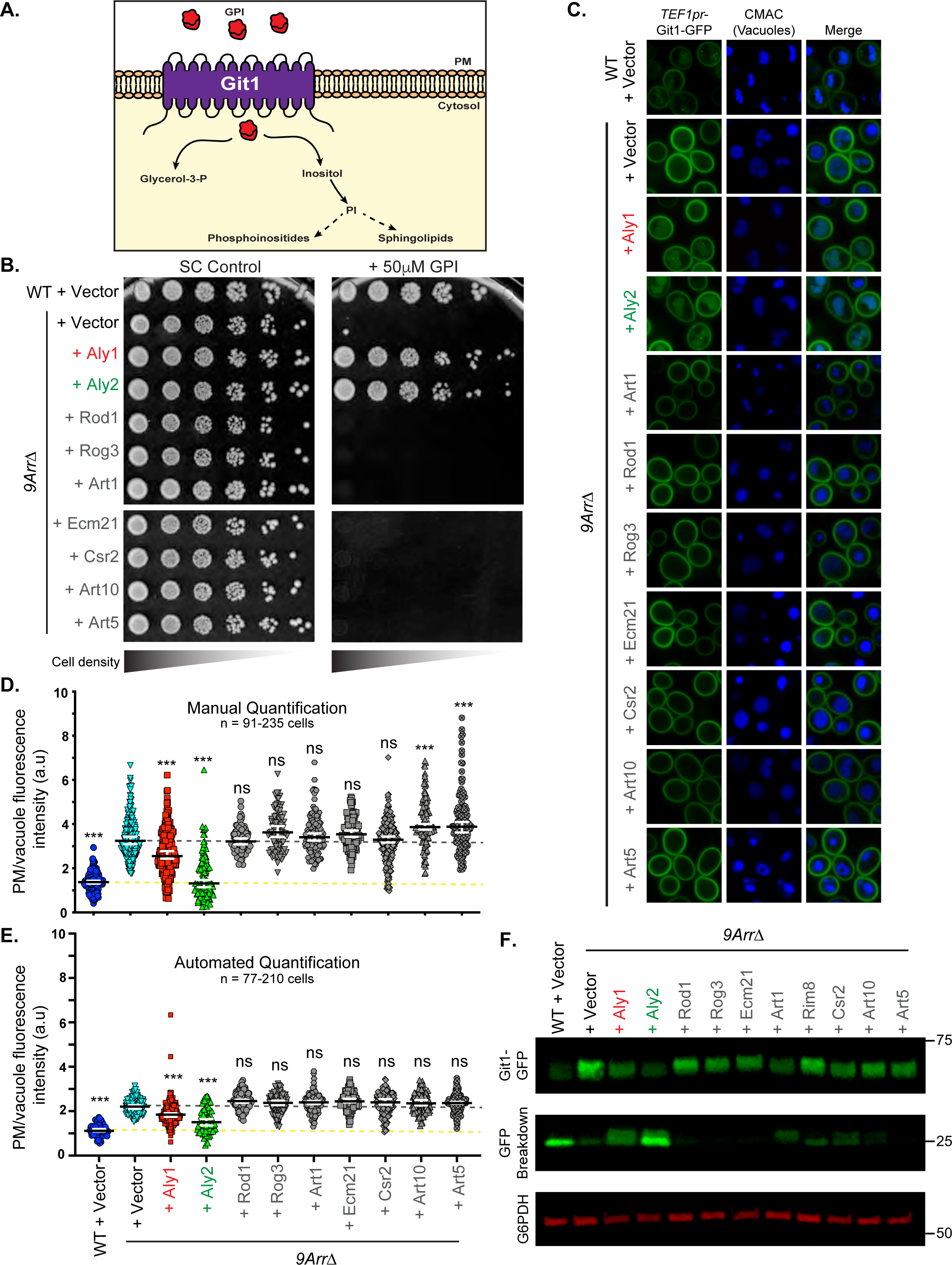
Loss of α-arrestins leads to retention of Git1 at the cell surface and sensitivity to GPI. *A*, Model of Git1 function at the plasma membrane and influence on downstream metabolic processes. Git1 primarily uptakes GPI, which is then converted to inositol and glycerol-3-phosphate. The inositol can feed into phosphatidylinositol (PI) synthesis, and this influences phosphoinositides and sphingolipids in the cell. *B*, Growth of serial dilutions of WT or *9ArrΔ* cells all containing a *LEU2*-marked *TEF1pr*-*GIT1*-GFP plasmid and the indicated pRS316-derived plasmids expressing either nothing (vector) or the indicated α-arrestins on SC medium lacking uracil and leucine and containing 50μM GPI. Growth is shown at 2 days after incubation at 30°C. *C*, WT or *9ArrΔ* cells containing Git1-GFP expressed from the *TEF1* promoter and either a pRS316 empty vector or plasmid expressing the indicated α-arrestins were imaged by confocal fluorescence microscopy. *D and E*, The PM and vacuolar fluorescence intensities from the cells depicted in C were quantified using either our manual analyses (D) or automated quantification (E) pipelines and the distributions of PM/intracellular fluorescence ratios in arbitrary units (a.u.) were plotted as scatter plots. The horizontal midline in black represents the median and the 95% confidence interval is represented by the white error bars. A yellow dashed line is added as a reference for the median ratio for wild-type cells and the grey dashed line provides a reference for the median value for the 9*Arr*Δ cells. Kruskal-Wallis statistical analyses with Dunn’s post hoc test were performed, and statistical significance compared to the *9Arr*Δ strain containing vector control is presented. *F*, Whole cell extracts from cells expressing Git1-GFP and α-arrestin-containing plasmids were analyzed by SDS-PAGE and immunoblotting. Git1-GFP and the GFP-breakdown product abundance is assessed with an anti-GFP antibody and the anti-Zwf1 (G6PDH) antibody is used as a loading control. Molecular weights of standards are shown at right measured in kilodaltons.

Given that GPI transport via Git1 impacts PI synthesis and, in turn, important downstream biosynthetic pathways, it is critical to define the factors that govern this protein’s stability and/or trafficking. Git1 is a 12-pass transmembrane domain protein that functions at the PM and is a member of the Major Facilitator Superfamily (MFS) (Mariggio *et al*., 2006). It contains sugar transporter-like domains, suggestive of a shared structural relationship to the glucose transporters in *Saccharomyces cerevisiae*. While there is no clearly defined mammalian ortholog of Git1, it has been shown that the mammalian glucose transporter Glut2 can uptake GPI and is considered functionally analogous (Mariggio *et al*., 2006). The trafficking of the other transporters needed for inositol uptake, such as Itr1 and Itr2, and members of the related glucose transporter family, including Hxt1, Hxt3, and Hxt6, are each controlled by a family of protein trafficking adaptors known as the α-arrestins (Nikko and Pelham, 2009; Harada *et al*., 2010; O’Donnell *et al*., 2015; Hovsepian *et al*., 2017; Ivashov *et al*., 2020). We therefore sought to determine if the α-arrestins might similarly play a role in Git1 transporter trafficking, whose removal from the cell surface was previously linked to clathrin-mediated endocytosis and subsequent vacuolar degradation (Almaguer *et al*., 2006).

The α-arrestins, conserved from yeast to man, act as bridges between membrane proteins (referred to as cargos) and Rsp5, a ubiquitin ligase known to regulate the endocytosis of many nutrient permeases; the ortholog of which is Nedd4-2 (Rotin and Kumar, 2009). Internalization of many nutrient transporters, such as the Fur4 uracil permease, and the Mup1 methionine permease, can be triggered by the presence of excess substrate (i.e., uracil or methionine, respectively) and subsequent ubiquitination of these transporters by Rsp5 ensures their interaction with the endocytic machinery, many components of which have ubiquitin-interaction motifs (Blondel *et al*., 2004; Lin *et al*., 2008; Nikko and Pelham, 2009; Ivashov *et al*., 2020). Modification of cargos by even a single ubiquitin is often sufficient to induce internalization (Shih *et al*., 2000; Raiborg *et al*., 2002; Stringer and Piper, 2011). However, many of the cargos controlled in this manner lack the motifs to directly recruit Rsp5. Instead, the α-arrestins selectively bind to membrane cargos and simultaneously interact with the WW-domains of Rsp5 via their PPxY motifs (Gupta *et al*., 2007; Lin *et al*., 2008; Nikko *et al*., 2008; Becuwe *et al*., 2012; O’Donnell *et al*., 2013; Alvaro *et al*., 2014). In this way, the α-arrestins link the cargo destined for internalization to the machinery needed to control selective endocytosis of a widening array of membrane proteins (reviewed in this issue of Biology of the Cell (Kahlhofer *et al*., 2020)). α-Arrestin function is often regulated by phosphorylation, with dephosphorylation typically stimulating α-arrestin-mediated endocytosis and phosphorylation typically impairing endocytosis (O’Donnell, 2012; O’Donnell and Schmidt, 2019; Kahlhofer *et al*., 2020). As a result of this regulation, the cell is not continuously internalizing transporters that should be maintained at the cell surface, but only stimulates selective removal in response to the appropriate nutrient cues and signaling pathways as α-arrestins become modified to allow for trafficking.

Of significant relevance to our studies, α-arrestins Art5 controls inositol influx into cells by regulating the inositol-stimulated endocytosis of inositol transporter Itr1, and perhaps Itr2 (Nikko and Pelham, 2009; Yamagami *et al*., 2015). Alterations in cellular inositol balance, as occurs in response to mis-regulation of Itr1 due to altered Art5 activity, has broad reaching implications in the cell, including causing changes to mitochondrial cardiolipin production by altering CDP-diacylglycerol levels (Harada *et al*., 2010; Tamura *et al*., 2013) and restoring polarized trafficking to cells where phospholipid flippases are mutated (Yamagami *et al*., 2015). Thus, clear ties between inositol balance, phospholipids and α-arrestins are already established.

Here we add new players in regulating GPI and inositol balance in cells; We show that the paralogous *<u>a</u>rrestin-like <u>y</u>east proteins 1* and *2* (Aly1 and Aly2) act as endocytic adaptors for Git1. This is a novel membrane protein cargo for these α-arrestins, which have previously been demonstrated to regulate the trafficking of Gap1, Dip5 and Put4, three amino acid permeases, the G-protein coupled receptor, Ste3, an arsenite transporter, Arr3 (also known as Acr3), and the P-type ATPase involved in salt and lithium homeostasis, Ena1 (Hatakeyama *et al*., 2010; O’Donnell *et al*., 2010; O’Donnell *et al*., 2013; Crapeau *et al*., 2014; Hager *et al*., 2018; Wawrzycka *et al*., 2019; Nishimura *et al*., 2020; Sen *et al*., 2020). Though only a few transporters are clearly linked to Aly1 and Aly2 to date, the diversity of transmembrane proteins regulated by this family of trafficking adaptor and the signaling cues that dictate their function is rapidly expanding. In the case of Git1, we show that α-arrestins operate in basal, unstimulated conditions to control the steady-state turnover of Git1 and are also induced to internalize Git1 in the presence of excess substrate, GPI. Like many other transported molecules, we show that excessive uptake of GPI is toxic to cells, though the mechanism of this toxicity is currently unclear. Consistent with earlier studies, we find that Git1 endocytosis occurs via the clathrin-mediated pathway (Almaguer *et al*., 2006); however, there is likely an intracellular sorting pathway at work in regulating Git1 as well since endocytic blocks do not fully recapitulate the defective trafficking observed with loss of the Alys. Throughout this study we support our fluorescent imaging of Git1, used to monitor its localization changes, with quantification and report the development of an automated pipeline for quantifying both plasma membrane and vacuolar fluorescent signal abundances that we call ‘*CellQuant*’ (available from www.odonnelllab.com). We compare the results from this automated quantification to the gold standard of manual image quantification and find that it accurately recapitulates the results from our manual analyses.

While Aly1 and Aly2 share functional similarities, Aly2 appears to play a dominant role in trafficking of Git1, which is especially evident in basal turnover of Git1. Interestingly, while α-arrestin Aly1 is regulated by the protein phosphatase calcineurin (CN) and dephosphorylation of Aly1 by CN stimulates Aly1-mediated endocytosis of the aspartic acid transporter Dip5 (O’Donnell *et al*., 2013), when we explored CN-regulation of Aly1-controlled steady state turnover of Git1 we found only modest contributions of this signaling pathway. However, when we explored the GPI ligand-induced phenotypes we found that a phosphomimetic allele of Aly1, where serines and threonines at CN-regulated sites were mutated to glutamic acid, failed to rescue *aly1*Δ *aly2*Δ sensitivity to GPI and the mutant allele that mimicked the dephosphorylated state of Aly1 was better able to control GPI-induced Git1 internalization. Thus, CN appears to play a role in controlling the ligand-induced trafficking of Git1. In addition, we identify genetic interactions between the Alys and CN in the regulation of cell sensitivity to drugs that perturb sphingolipid biosynthesis. However, the phenotypic links to sphingolipid metabolism run counter to the increased sphingolipid flux that should arise when PI levels are elevated, and instead suggest a more complex regulatory connection between these players.

Finally, loss of the Alys results in a sharp shift in phosphatidyl-inositol-3-phosphate (PI(3)P) balance in the endo-membrane system, with a dramatic increase in PI(3)P levels in the limiting membrane of the vacuole where normally this lipid is largely sequestered to endosomes (Schu *et al*., 1993; Burd and Emr, 1998; Gillooly *et al*., 2000). This alteration in PI(3)P does not appear to reflect an overall alteration PIPs as other phosphoinositides (including phospho-inositol-4-phosphate (PI(4)P) and phospho-inositol-4,5-bisphosphate (PI(4,5)P2) are largely unchanged in the absence of the Alys. While uptake of GPI exacerbates this defect, with even more inappropriate PI(3)P retention on the vacuolar membrane in both wild-type and *aly1*Δ *aly2*Δ cells, it is unlikely that this change in PI(3)P abundance in *aly1*Δ *aly2*Δ is entirely due to GPI uptake via Git1. However, we confirm that [^3^H]inositol derived from exogenously supplied [^3^H]-inositol-GPI is incorporated into PI (Patton *et al*., 1995; Patton-Vogt and Henry, 1998), the precursor to both the inositol-containing complex sphingolipids and PIPs. Here we further show that this label incorporation into PI increases in the absence of Aly and Aly2. Thus overall, we show that the α-arrestins Aly1 and Aly2 are new regulators of GPI uptake via Git1 and, thereby, can impact cellular PI levels and downstream lipid species.

## Results

### Trafficking of Git1 is impaired in the absence of α-arrestins Aly1 and Aly2

We sought to determine if the α-arrestins are involved in controlling Git1 localization. Initially, in order to avoid confounding issues linked to endogenous *GIT1* gene expression, we expressed Git1-GFP from the constitutive *TEF1* promoter and examined growth of cells lacking 9 of the 14 yeast α-arrestins (referred to hereafter as 9ArrΔ strain (Nikko and Pelham, 2009)) on medium containing Git1’s substrate, GPI. We found that 9ArrΔ cells were exquisitely sensitive to growth on medium containing GPI, while wild-type cells grew well at the concentrations of GPI employed (Figure 1B). Interestingly, functional complementation assays where plasmids expressing each of the missing α-arrestins in the 9ArrΔ cells demonstrated that addition of either Aly1, or its paralog Aly2, restored growth of 9ArrΔ cells to near wild-type levels in the presence of GPI. We hypothesized that the sensitivity to GPI displayed by the 9ArrΔ cells might arise due to excessive GPI uptake and imbalance of downstream metabolites and pathways (Figure 1A). To determine if Git1 was retained at the cell surface in 9ArrΔ cells, we assessed its steady-state localization using confocal microscopy. Consistent with potentially elevated GPI import in the 9ArrΔ cells, we found that Git1 was dramatically retained at the cell surface in these mutant cells while in wild-type cells there was a balance of Git1 localized faintly to the plasma membrane with a substantial amount of the fluorescent signal in the vacuole (Figure 1C). Additionally, the retention of Git1 at the plasma membrane in 9ArrΔ cells was diminished in cells expressing either *ALY1* or *ALY2*. However, plasmids expressing the other α-arrestins failed to improve growth on GPI or reduce the plasma membrane retention of Git1 at the cell surface (Figures 1B-C). Manual quantification of the plasma membrane (PM) and vacuolar fluorescence intensities for Git1-GFP supported these conclusions, showing significantly reduced PM/vacuole fluorescence ratios in wild-type cells compared to the 9ArrΔ cells (Figure 1D). However, addition of either Aly1 or Aly2 significantly reduced the PM/vacuole ratio of 9ArrΔ cells, with Aly2 having the more robust effect (Figure 1D). Addition of the other α-arrestins to 9ArrΔ cells did not significantly reduce the PM/vacuole fluorescence ratio, yet some did significantly increase the PM/vacuole ratio, suggesting that their expression might further impair Git1 internalization or trafficking to the vacuole in the 9ArrΔ cells.

Our lab has made extensive use of this kind of manual fluorescence quantification (O’Donnell *et al*., 2015; Prosser *et al*., 2015; Chandrashekarappa *et al*., 2016; Hager *et al*., 2018; Soncini *et al*., 2020) and the process is time consuming and limits the number of cells that can be assessed; For manual quantification, we typically quantify at least 150 cells with at least two full imaging fields of the cells being measured (O’Donnell *et al*., 2015). To facilitate this quantification, we developed an automated pipeline for this process using an R-script program we wrote called ‘*CellQuant*’ (see Materials and Methods; script available at www.odonelllab.com and on Github at https://github.com/sah129/CellQuant). We re-ran our analyses to measure the PM and vacuolar fluorescence intensity for the images in Figure 1C using our automated pipeline and had strong agreement with our manual data set. The time taken to obtain this quantification was a small fraction of that required for the manual quantification (∼30 mins compared to >20 hours) (Figure 1E). The imaging data, both types of quantification, and GPI growth assays were consistent with the results of our immunoblot analyses of Git1-GFP abundance, where we found much less Git1-GFP in protein extracts from wild-type cells than in those from 9ArrΔ cells. In wild-type cells there was a significant signal observed for the free GFP breakdown product, often associated with the stabilized cleaved GFP in the vacuole after partial degradation of GFP-tagged transporters (Li *et al*., 1999; Nikko and Pelham, 2009), while in the 9ArrΔ cells there was very little signal for the GFP-breakdown product (Figure 1F). Addition of either Aly1 or Aly2 to 9ArrΔ cells reduced the Git1-GFP total protein abundance in protein extracts dramatically and increased the GFP-breakdown product associated with vacuolar degradation of the transporter (Figure 1F). Expression of the other α-arrestins in the 9ArrΔ cells had modest effects on total Git1-GFP levels and did very little to increase the abundance of the GFP breakdown product (Figure 1F). Taken together these data show that the α-arrestins Aly1 and Aly2 play an important role in trafficking of Git1 to the vacuole for degradation, either by stimulating its endocytosis or its intracellular sorting from the Golgi and/or endosomes to the vacuole. Additionally, these findings validate that our new automated quantification pipeline is effective for measuring PM and vacuole fluorescence abundance and we will continue to assess its utility in the coming analyses throughout this paper.

Our findings to date have hinged on the use of the 9ArrΔ cells, however this background is missing only 9 of the 14 yeast α-arrestins. Given the data indicating that addition of some of the α-arrestins to 9ArrΔ cells resulted in increased PM retention of Git1, we sought to determine if the other α-arrestins that are intact in 9ArrΔ cells might contribute to Git1 trafficking. A recent paper demonstrated that the α-arrestins and other adaptors that bind the ubiquitin ligase Rsp5 may compete for a limited pool of Rsp5 and therefore restrict its function (MacDonald *et al*., 2020). We observed that when α-arrestins other than Aly1 or Aly2 were expressed in the 9ArrΔ background there were elevated Git1 levels at the cell surface (Figure 1C-D). This result might be obtained if the expression of an α-arrestin that was unable to traffic Git1 somehow interferes with the activity of one of the remaining endogenous α-arrestin adaptors. In addition, the 9ArrΔ cells still retain 5 other α-arrestins, including Bul1 and Bul2, which have overlapping functions in other Aly1/Aly2-mediated trafficking events (Novoselova *et al*., 2012; Crapeau *et al*., 2014), and so we next determined if the Bul proteins might also influence Git1 trafficking. Interestingly, growth assays on GPI did not show any significant change in sensitivity of the *bul1*Δ *bul2*Δ cells relative to wild-type cells, however loss of just the *bul2*Δ on its own gave limited growth at higher concentrations of GPI (Supplemental Figure 1A). When we constructed deletions of *aly1*Δ *aly2*Δ and either one or both of the *BUL*s we found that cells lacking *BUL2* in addition to *ALY1 and ALY2* were more sensitive to GPI (Supplemental Figure 1A; see the 25μM GPI growth assays). These data suggest that Bul2, but not Bul1, may also play a role in regulating Git1 trafficking under some circumstances. Furthermore, imaging of Git1 in the absence of Bul proteins displayed a surprising pattern of localization with greatly reduced Git1 abundance in all cells lacking any combination of Bul proteins (Supplemental Figure 1B-C). Perhaps this is an indication that Bul proteins influence expression from the *TEF1* promoter used for *GIT1* in these experiments, though to date all of the ascribed effects of α-arrestins have been post-transcriptional in nature. Given the complex results with these mutants, the contributions of Buls to Git1 trafficking will be explored in more detail elsewhere, and here we focus our experimental attention on the roles of Aly1 and Aly2.

To confirm that the paralogous α-arrestins Aly1 and Aly2 were important for Git1 trafficking, we assessed the sensitivity of *aly1*Δ *aly2*Δ cells to GPI in comparison to wild-type and 9ArrΔ cells. We found that loss of either *ALY1* or *ALY2* sensitized cells to GPI, with loss of *ALY2* having a more severe growth phenotype (Figure 2A). Consistent with a role for Aly1, cells lacking both *ALY1* and *ALY2* were more sensitive to GPI than either single mutant and had the same degree of GPI sensitivity as the 9ArrΔ cells (Figure 2A). Fluorescent imaging of Git1-GFP in *aly1*Δ *aly2*Δ cells revealed Git1 retention at the PM and significantly increased PM/vacuole ratio in these cells that is comparable to that of 9ArrΔ, as judged by both our manual and automated quantification pipelines (Figures 2B-D). Loss of *ALY2* alone dramatically increased Git1-GFP at the PM, while deletion of *ALY1* had no significant impact, suggesting that under these steady-state conditions Aly1 has little impact on Git1 trafficking (Figures 2B-D).

**FIGURE 2.**
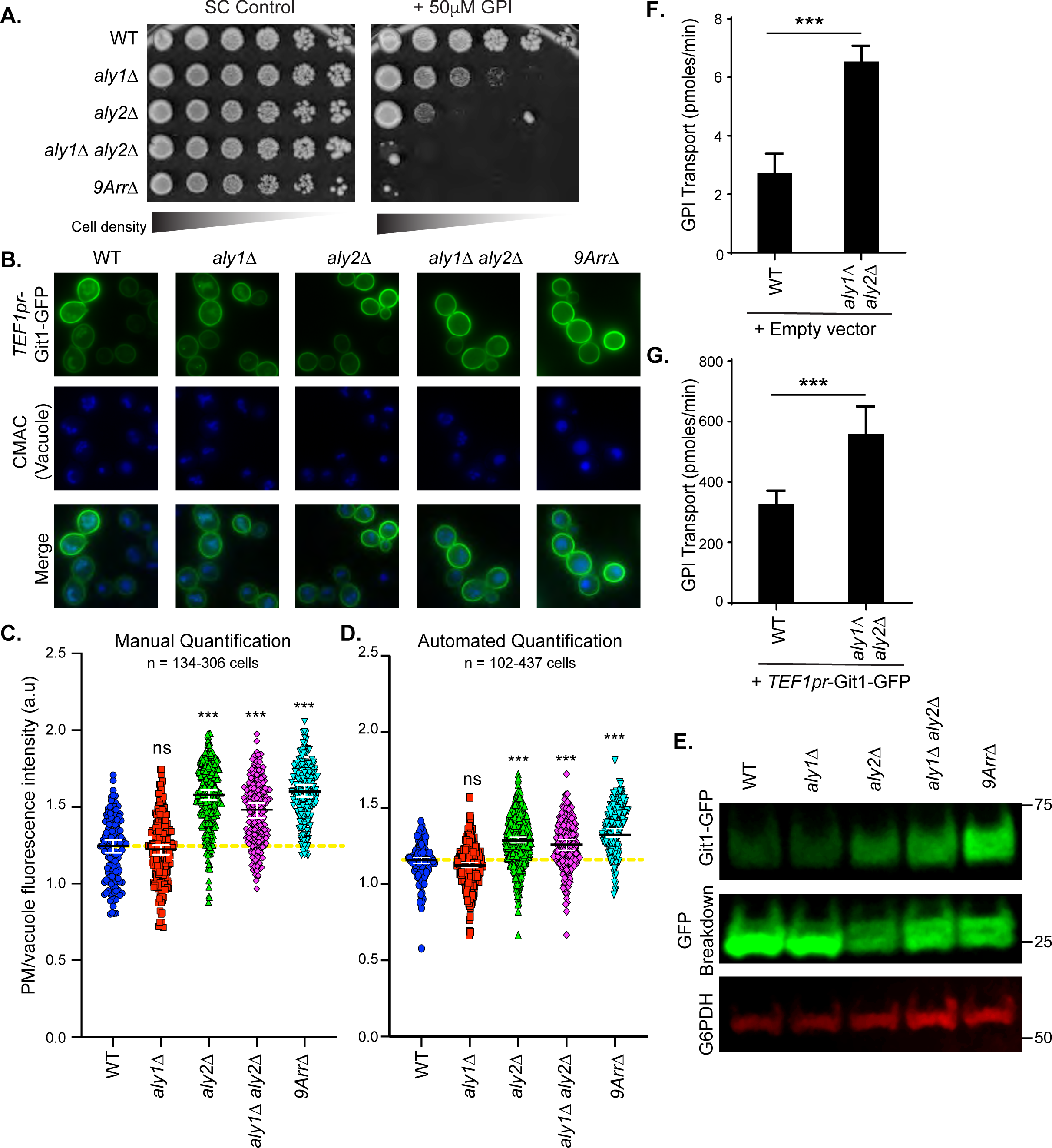
Aly1 and Aly2 regulate Git1 localization at the cell surface and loss of these α-arrestins causes GPI sensitivity. *A*, Growth of serial dilutions of WT, *aly1Δ, aly2Δ, aly1Δ aly2Δ*, or *9ArrΔ* cells containing the *TEF1pr*-*GIT1*-GFP plasmid on SC medium lacking leucine and containing 50μM GPI. Growth is shown at 2 days after incubation at 30°C. *B*, The indicated cells containing Git1-GFP expressed from the *TEF1* promoter were stained with CMAC and imaged by fluorescence microscopy. *C and D*, The PM and vacuolar fluorescence intensities from the cells depicted in B were quantified using either our manual (C) or automated quantification (D) pipelines and the distributions of PM/vacuolar fluorescence ratios in arbitrary units (a.u.) were plotted as scatter plots. The horizontal midline in black represents the median and the 95% confidence interval is represented by the white error bars. A yellow dashed line is added as a reference for the median ratio for wild-type cells. Kruskal-Wallis statistical analyses with Dunn’s post hoc test were performed, and statistical significance compared to the wild-type cells is presented. *E*, Whole cell extracts from the indicated cells containing Git1-GFP expressed from the *TEF1* promoter were resolved by SDS-PAGE and immunoblotted with an anti-GFP antibody (to detect Git1-GFP and the GFP breakdown product) and anti-Zwf1 antibody (G6PDH) as a loading control. Molecular weights of standards are shown at right measured in kilodaltons. *F and G,* GPI radio-labelled uptake assays were performed as described in Surlow, et. al. 2014. WT and *aly1Δ aly2Δ* cells containing a *LEU2*-marked *TEF1pr*-GFP empty vector (*F)* or a *LEU2*-marked *TEF1pr*-*GIT1*-GFP plasmid *(G)* were grown at 30°C to logarithmic phase in synthetic media lacking inositol and choline and containing 10mM inorganic phosphate. 50μL of 25μM of [^3^H]GPI *(F)* or 75μM of [^3^H]GPI *(G)* was added to 200μL of processed cell suspension. Suspension was then filtered and washed before resuspending in 10mL of scintillation fluid for counting. Statistical t-test value represents means ± SD of biological triplicates performed in duplicate (6 replicates in total).

Consistent with the idea that Aly2 has a larger influence on Git1 trafficking in steady state conditions, protein extracts from *aly2*Δ or *aly1*Δ *aly2*Δ cells revealed little difference between Git1-GFP abundance, and both of these strains had higher levels of Git1-GFP than those from either *aly1*Δ or wild-type cells (Figure 2E). In 9ArrΔ cells, the Git1 levels appear marginally higher than in the *aly1*Δ *aly2*Δ cells, in line with the fluorescence imaging data (Figures 2B-E). It should be noted that while none of the other α-arrestins tested in this background reduced Git1 levels (Figure 1 B-F), this strain is additionally missing Bsd2, a known factor in sorting membrane proteins in the Golgi. Therefore, *BSD2* loss could be making a modest contribution to Git1 sorting in 9ArrΔ cells explaining the altered abundance of Git1 in 9ArrΔ compared to *aly1*Δ *aly2*Δ cells.

To this point, our assays used a plasmid-borne version of Git1 that was GFP tagged. To confirm that this version of Git1 was functional and accurately reflected events that regulate endogenous Git1, we performed short-term radiolabeled GPI transport assays (5 min) in wild-type and *aly1*Δ *aly2*Δ cells with and without the Git1-GFP plasmid. These cells were grown in high phosphate medium, where expression of Git1 would be at basal levels and not induced as it would be under phosphate or inositol starvation conditions (Almaguer *et al*., 2003). Even under these high phosphate conditions, we found that loss of α-arrestins *ALY1* and *ALY2* increased the rate of GPI transport into cells by more than ∼3-fold compared to wild-type cells (Figure 2F). To ensure that the Git1-GFP construct was functional, we also assessed radiolabeled GPI uptake in wild type or *aly1*Δ *aly2*Δ cells harboring the Git1-GFP plasmid. In cells expressing Git1-GFP from a constitutive promoter, GPI uptake was increased by ∼100-fold in wild-type cells grown in high phosphate medium (compare Figures 2F and 2G). Cells lacking *ALY1* and *ALY2* nearly doubled this rate of uptake (Figure 2G), again consistent with retention of Git1 at the cell surface in the absence of these α-arrestins. In addition to these uptake assays, we examined the ability of the Git1-GFP plasmid to rescue growth of *git1*Δ on medium lacking phosphate and containing GPI. Earlier studies found that cells can grow when supplemented with GPI as sole phosphate source (Almaguer *et al*., 2004). We show that growth on media containing GPI a sole phosphate source was rescued to wild-type levels in *git1*Δ cells expressing the Git1-GFP plasmid (Supplemental Figure 2A). These data also demonstrate that the GFP-tagged Git1 is functional, as its expression increases GPI transport into cells, even when the endogenous Git1 was not expressed.

To more carefully assess regulation of endogenous Git1, we made a chromosomally integrated, GFP-tagged version of this gene under its own promoter. We then examined Git1-GFP localization in cells grown in medium that contained high or low levels of phosphate, with or without inositol. Expression of Git1 has previously been shown to be upregulated in response to phosphate and inositol limitation as a result of the *GIT1* promoter’s regulation by Pho2/Pho4 and Ino2/Ino4 transcription factors under these respective conditions (Almaguer *et al*., 2003). As expected, regardless of the inositol content, in high phosphate medium there was little to no detectible Git1 by fluorescence microscopy (Figures 3A-B, note that 9ArrΔ cells in these conditions were sick and faint GFP fluorescence observed for one cell was from a dead cell). However, when these cells were incubated in low phosphate medium Git1-GFP fluorescence was observed (Figures 3C-D). Under low phosphate conditions, the wild-type cells had predominantly vacuolar GFP fluorescence, yet *aly1*Δ *aly2*Δ or the 9*Arr*Δ cells retained Git1-GFP at the PM. The fluorescence intensity for each of these was slightly brighter when cells were additionally incubated in medium without inositol, consistent with the earlier studies of *GIT1* expression (Almaguer *et al*., 2003). We performed manual quantification on this data set and find that there is substantially brighter PM/vacuole fluorescence ratio in the *aly1*Δ *aly2*Δ and 9*Arr*Δ cells in low phosphate medium with or without inositol (Figures 3E-F). We also performed automated quantification of these data sets but used a slightly different algorithm that made use of the DIC channel image to define the cell surface since the PM fluorescence for the wild-type cells was too low and made it difficult to capture using our standard approach (see Methods). These data agree with the manual quantification we performed, although there was some compaction in the range of values identified in the automated quantification compared to the manual version (Supplemental Figures 2B-C). Taken together, these findings are consistent with our observations that the α-arrestins, in particular Aly1 and Aly2, contribute to Git1 trafficking and their absence results in retention of Git1 at the cell surface.

**FIGURE 3.**
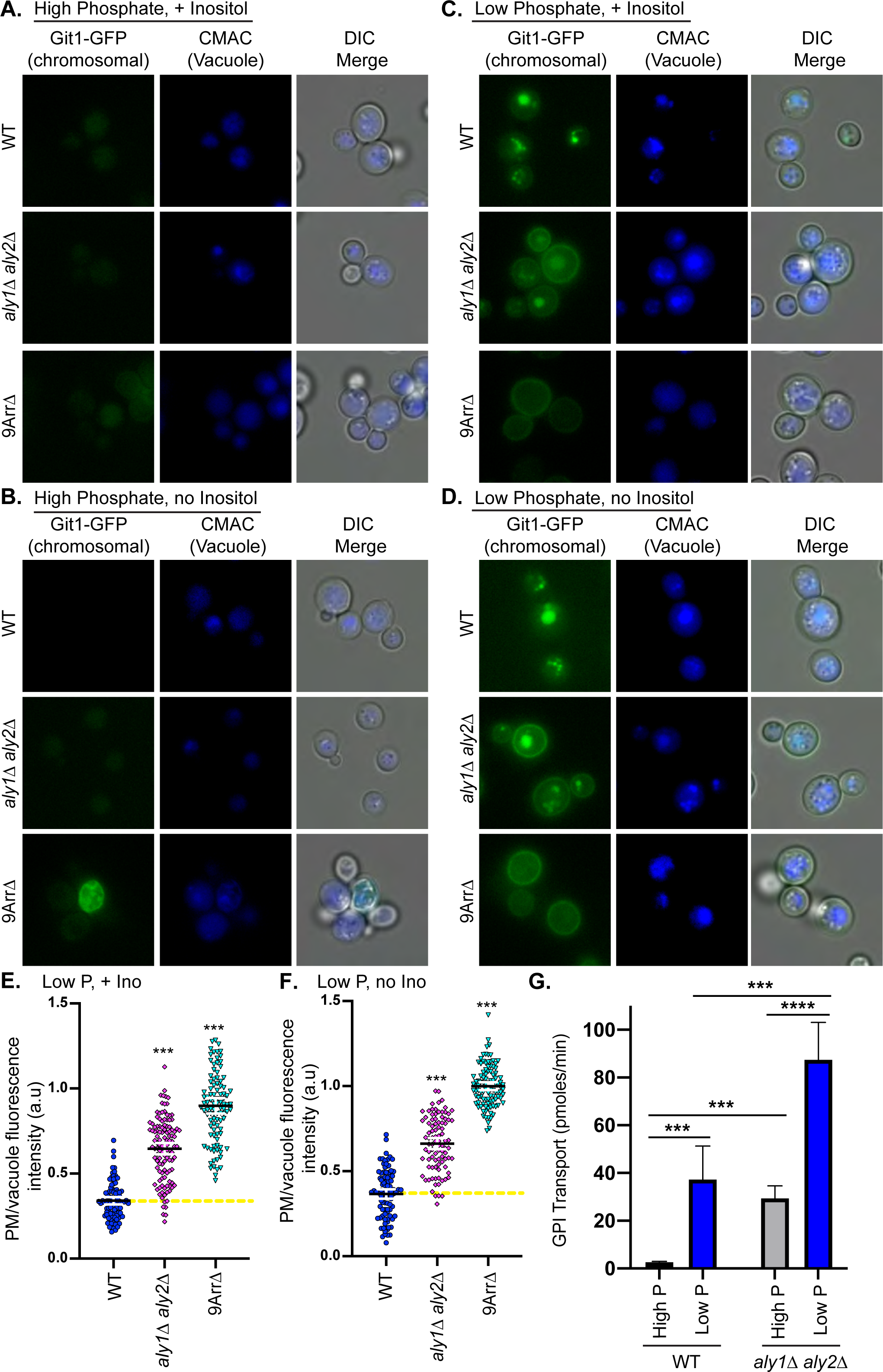
Aly1 and Aly2 regulate endogenous Git1 transporter localization and function. *A-D*, The indicated cells containing a chromosomally integrated Git1-GFP at its endogenous locus were grown in medium containing high (10mM) or low (100μM) phosphate with or without inositol for 12 h, stained with CMAC and imaged using a fluorescence microscope. *E and F*, The images presented in panels C and D were quantified using our manual quantification methodology and the distributions of PM/ vacuolar fluorescence ratios in arbitrary units (a.u.) were plotted as scatter plots. The horizontal midline in black represents the median and the 95% confidence interval is represented by the white error bars. A yellow dashed line is added as a reference for the median ratio for wild-type cells. Kruskal-Wallis statistical analyses with Dunn’s post hoc test were performed and statistical significance compared to wild-type cells is presented. *G*, GPI radio-labelled uptake assays were performed as described in Surlow, et. al. 2014. WT and *aly1Δ aly2Δ* cells were grown at 30°C to logarithmic phase in synthetic media lacking inositol and choline and containing 100μM (low) or 10mM (high) inorganic phosphate. [^3^H]GPI was added to 200μL of processed cell suspension for 5 min and cells were filtered and washed before resuspending in 10mL of scintillation fluid for counting. Statistical t-test value represents means ± SD of biological triplicates performed in duplicate (6 replicates in total).

We further explored the role of Aly1 and Aly2 in regulating Git1 function under shifting phosphate conditions using a short-term GPI transport assays (Surlow *et al*., 2014). When wild-type cells were shifted from high to low phosphate medium to induce Git1 expression, GPI transport increased ∼14-fold (Figure 3G). In *aly1*Δ *aly2*Δ cells grown in low phosphate medium, the GPI transport rate was ∼3-fold higher than that of wild-type cells in low phosphate or *aly1*Δ *aly2*Δ cells in high phosphate medium (Figure 3G). These data, taken together, demonstrate that the loss of *ALY1* and *ALY2* increases GPI uptake by endogenous, untagged Git1 in response to phosphate starvation and are consistent with our findings that Git1-GFP is retained at the cell surface in *aly1*Δ *aly2*Δ cells (Figures 2B-D, 3C-F and Supplemental Figure 2B-C).

### Git1 internalization is induced by GPI and is dependent upon α-arrestins Aly1 and Aly2

Our experiments to this point indicate that Aly1 and Aly2 play a role in the steady-state internalization and/or trafficking to the vacuole of Git1, with Aly2 performing the vast majority of this trafficking and Aly1 only showing a dramatic alteration of Git1 localization when present in the 9ArrΔ cells (Figure 1B-F). However, for many transporters, addition of their substrate triggers conformational changes that can stimulate their sorting to the vacuole and/or endocytic turnover (Gournas *et al*., 2010; Keener and Babst, 2013; Ghaddar *et al*., 2014; Gournas *et al*., 2017). We monitored the distribution of Git1-GFP in response to GPI addition and found that Git1 fluorescence shifted to the lumen of the vacuole in wild-type cells after GPI addition (Figures 4A-C). This GPI-induced shift in Git1 localization was largely lost in *aly1*Δ *aly2*Δ cells, though some modest reduction in PM/vacuole ratio is observed at 2h post GPI addition, suggesting a GPI-induced and Aly1/Aly2-independent mechanism of reducing Git1 abundance during prolonged GPI exposure (Figures 4A-C). Introduction of plasmid-borne Aly1 to *aly1*Δ *aly2*Δ cells significantly improved the GPI-induced trafficking to the vacuole for Git1 (Figures 4A-C; compare 60 min treatment between Aly1 vs vector control). Interestingly, the ratio of PM/vacuole intensity in *aly1*Δ *aly2*Δ cells is initially still significantly higher than in wild-type cells when Aly1 is present, consistent with our earlier data showing that Aly1 has a minor role to play in the steady-state trafficking of Git1 to the vacuole (Figures 4A-C and Figure 2). Addition of Aly2 to *aly1*Δ *aly2*Δ cells was better able to restore basal trafficking of Git1 to the vacuole, as indicated by the reduced PM/vacuole fluorescence ratios observed prior to GPI addition, which was not significantly different from the PM/vacuole ratios in wild-type cells (Figures 4A-C). However, GPI-induced trafficking was not as impacted by the presence of Aly2, with only modest drops to the PM/vacuole ratios at 60- and 120-mins post GPI addition, similar to what was observed for *aly1*Δ *aly2*Δ cells with just a vector control. Importantly, our manual and automated quantification of this imaging (Figures 4B and 4C, respectively) are consistent with one another. Taken together, these data support a model whereby Aly1 may play a more prominent role in the GPI-induced trafficking of Git1, while Aly2 is very effective at steady-state trafficking of Git1 to the vacuole. These data further suggest that there may be additional players in this trafficking as there was some turnover of Git1 after extended incubations in GPI in the *aly1*Δ *aly2*Δ cells. Perhaps this could be ascribed to Bul2 function or another trafficking adaptor that can operate in this GPI-stimulated turnover pathway when the *ALY*s are deleted.

**FIGURE 4.**
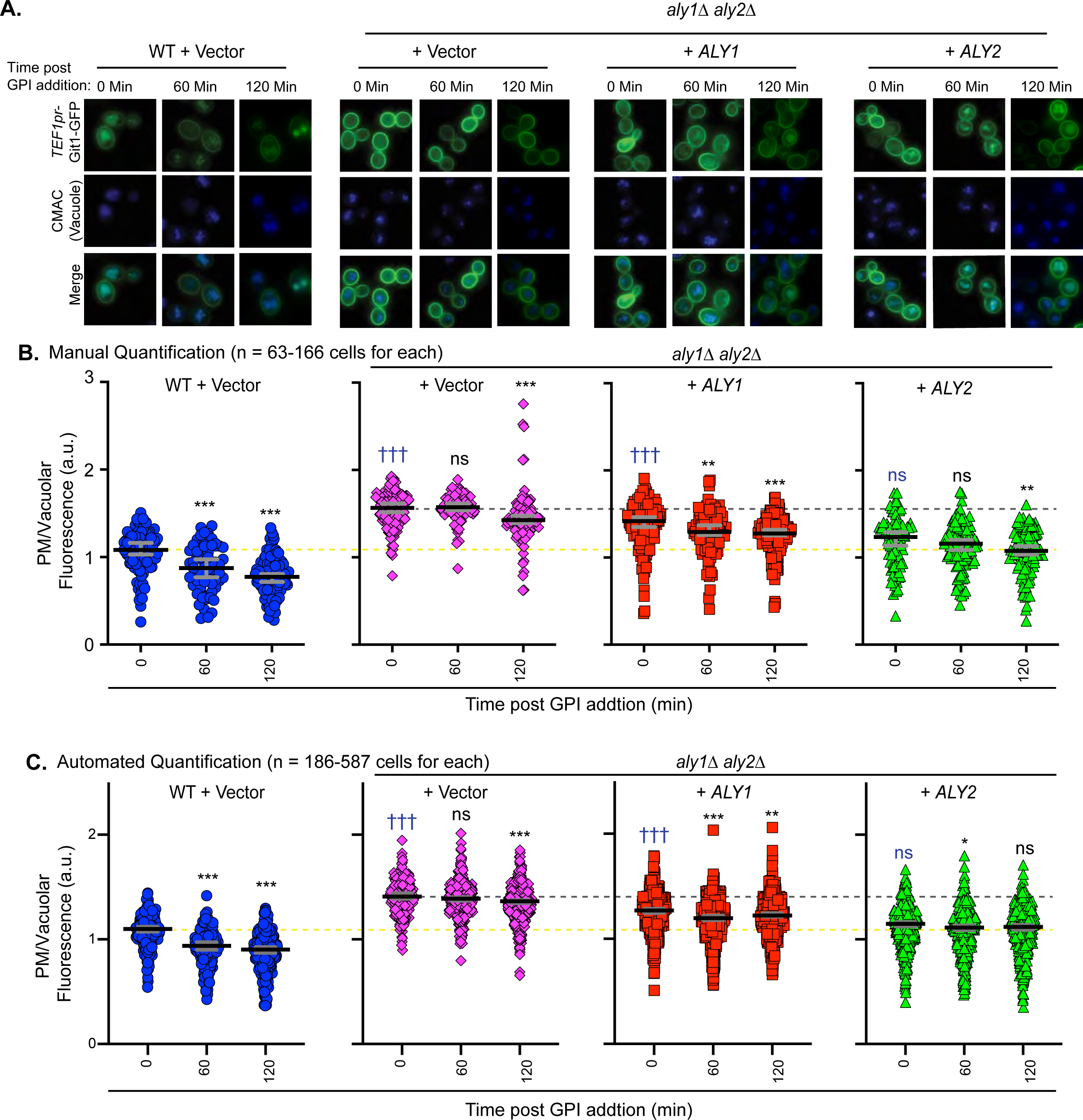
Git1 trafficking to the vacuole is induced by GPI addition and Aly1 and Aly2 regulate Git1 sorting to the vacuole. *A,* WT or *aly1Δ aly2Δ* cells containing Git1-GFP expressed from the *TEF1* promoter and either an pRS316 vector expressing nothing or one expressing the indicated α-arrestin were cultured in SC media lacking uracil and leucine and containing 50μM GPI for the times indicated before being imaged by fluorescence microscopy. *B and C*, The PM and vacuolar fluorescence intensities from the cells depicted in A were quantified using either our manual (B) or automated quantification (C) pipelines and the distributions of PM/vacuolar fluorescence ratios in arbitrary units (a.u.) were plotted as scatter plots. The horizontal midline in black represents the median and the 95% confidence interval is represented by the grey error bars. Kruskal-Wallis statistical analyses with Dunn’s post hoc test were performed, and statistical significance compared to the 0 min point for each graph is presented as asterisks (*) or indicated as not significant (ns). Additionally, the 0 min point for each of the strains is compared back to the wild-type with vector control and these are presented in blue text as daggers (†) or indicated as not significant (ns) where applicable. A yellow dashed line is presented to facilitate comparisons to the WT with vector control at the t=0 min point, and a grey dashed line is presented to facilitate comparisons to the *aly1*Δ *aly2*Δ mutant with vector at the t=0 min point.

To begin to define the relative contributions of endocytic trafficking versus intracellular sorting on Git1 accumulation in the vacuole under basal conditions and after GPI addition, we examined the Git1-GFP localization and GPI sensitivity of mutant cells lacking endocytic components. As clearly demonstrated in our earlier assays, loss of *ALY1* and *ALY2* resulted in exquisite sensitivity to GPI and retention of Git1-GFP at the cell surface (Figures 5A-C and 2A-C). Loss of the endocytic factor *END3*, which encodes an early arriving component of endocytic patches that helps in clathrin-mediated endocytosis, similarly caused sensitivity to GPI and retention of Git1-GFP at the PM (Figures 5A-C). Interestingly, deletion of *APL3* or *APL1*, which encode the α- and *β*-adaptin, respectively, of clathrin-associated protein complex 2 (AP-2), each gave rise to increased sensitivity to GPI, with *apl1*Δ cells having a more severe phenotype than *apl3*Δ cells (Figure 5A). While the role of AP-2 cargo selection for clathrin-mediated endocytosis is well-documented in mammals, in yeast this complex has only been associated with the internalization of a few membrane proteins, though it clearly associates with early endocytic patches (Carroll *et al*., 2009; Carroll *et al*., 2012). The rationale for examining AP-2 in these studies is because Aly1 and Aly2 have been shown to physically interact with these adaptins (O’Donnell *et al*., 2010). In line with the GPI sensitivity results, loss of *APL1* led to retention of Git1 at the cell surface, while *apl3*Δ significantly impaired vacuolar trafficking of Git1, but did not do so as robustly as *apl1*Δ (Figures 5B-C). Based on these data, we suggest that a portion of the steady-state trafficking of Git1 to the vacuole occurs via endocytosis, as there is reduced vacuole-localized GFP fluorescence in the endocytic mutants. These data further suggest that AP-2 may play a role in regulating Git1 endocytosis. However, it is interesting that the *aly1*Δ *aly2*Δ cells show a more robust retention of Git1 at the PM than any of these other mutants, indicating that the α-arrestins may additionally impair intracellular sorting of Git1.

**FIGURE 5.**
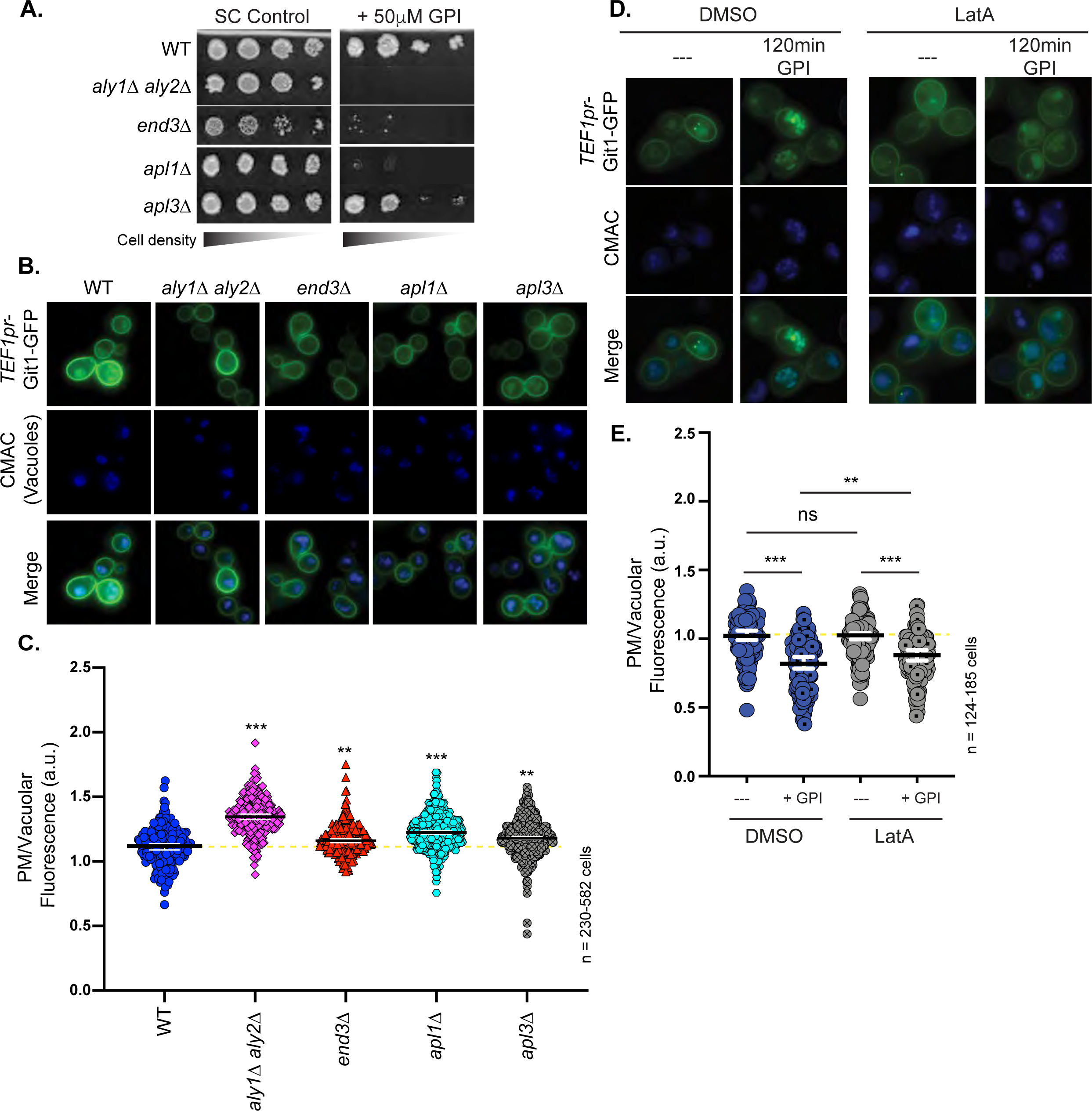
GPI induces clathrin-dependent endocytosis of Git1. *A,* Growth on SC medium lacking leucine with or without 50μM GPI for serial dilutions of cells from the indicated yeast strains containing a *LEU2*-marked *TEF1pr*-*GIT1*-GFP plasmid is shown 2 days post-incubation at 30°C. *B*, The indicated cells expressing Git1-GFP from the *TEF1* promoter were stained with CMAC and imaged by fluorescence microscopy. *C and E*, The PM and vacuolar fluorescence intensities from the cells depicted in B or D, respectively, were quantified using our automated quantification pipeline and the distributions of PM/intracellular fluorescence ratios in arbitrary units (a.u.) were plotted as scatter plots. The horizontal midline in black represents the median and the 95% confidence interval is represented by the white error bars. A yellow dashed line represents the median ratio for untreated wild-type cells to facilitate comparisons. Kruskal-Wallis statistical analyses with Dunn’s post hoc test were performed and statistical significance is indicated with asterisks. *D,* WT cells containing Git1-GFP expressed from the *TEF1* promoter were cultured in SC media lacking leucine and containing 200μM latrunculin A or an equal volume of DMSO vehicle for 120 min prior either addition of 50μM GPI or no treatment, and incubated for 120 min further before being stained with CMAC and imaged by fluorescence microscopy.

To examine ligand-induced internalization and intracellular trafficking, we next examined the localization of Git1-GFP in cells treated with GPI that had either been preincubated in <u>lat</u>runculin <u>A</u> (LatA), which prevents actin polymerization to selectively block endocytosis while leaving intracellular sorting intact (Ayscough *et al*., 1997), or its vehicle control DMSO. Wild-type cells treated with LatA retained higher PM/vacuole fluorescence ratios after GPI addition (Figure 5D-E). However, the ratio was still significantly higher than the vehicle control treated cells after 2h of GPI incubation, indicating that some of the trafficking to the vacuole was likely due to endocytosis while a portion of the trafficking was independent of this LatA blocked pathway. Interestingly, the LatA treated cells were not statistically different from the DMSO-treated cells prior to GPI addition, suggesting that a portion of Git1 likely travels to the vacuole through an intracellular sorting route (Figure 5D-E). For *aly1*Δ *aly2*Δ cells treated this same way, the GPI induced a small amount of Git1-GFP trafficking to the vacuole (as measured by decreased PM/vacuole fluorescence ratio) and Aly1 addition dropped this further but the effect was muted in the LatA treated cells (Supplemental Figure 3A). For the case of cells expressing Aly2 only, there was no impact of having GPI added if the cells were pretreated with LatA (Supplemental Figure 3A). Thus, although GPI induces endocytosis of Git1 and this accounts for a portion of the increased vacuolar signal observed, the persistent decrease in the PM/vacuole ratio observed in LatA treated cells after GPI addition suggests that there is a contribution of intracellular sorting to the vacuole for Git1-GFP in GPI treated cells. Aly1 appeared better able to stimulate this intracellular sorting, as LatA treated Aly1-expressing cells significantly dropped the PM/vacuole ratios after GPI treatment. In contrast, Aly2 did not seem to operate in this pathway as cells expressing Aly2 failed to reduce PM/vacuole ratios after LatA and GPI treatments.

### α-Arrestins Aly1 and Aly2 must interact with Rsp5 to regulate Git1 trafficking

α-Arrestins all contain ^L^/_P_PxY motifs which allow for association with WW-domain-containing proteins. In yeast, each of the α-arrestins has been shown to use these ^L^/_P_PxY motifs to engage the ubiquitin ligase Rsp5, which harbors three WW-domains (Figure 6A) (Gupta *et al*., 2007; Lin *et al*., 2008; O’Donnell *et al*., 2013; Becuwe and Leon, 2014). In addition to binding Rsp5, the α-arrestins bind to membrane proteins, thereby bringing the ubiquitin ligase in close proximity to many of its membrane client proteins that may lack their own ^L^/_P_PxY motifs. Ubiquitination of membrane proteins serves as a signal for their endocytosis, with many of the early-arriving endocytic proteins having ubiquitin-interaction motifs that aid in cargo clustering at endocytic sites (Weinberg and Drubin, 2012). Not only do the membrane cargo proteins get ubiquitinated by Rsp5, but many of the α-arrestins themselves are mono-ubiquitinated by this ubiquitin ligase (Gupta *et al*., 2007; Lin *et al*., 2008; Becuwe *et al*., 2012; O’Donnell *et al*., 2013; Ho *et al*., 2017). This mono-ubiquitination is thought to be activating for α-arrestin function, and in some cases is required for α-arrestin recruitment to the membrane (Lin *et al*., 2008; MacGurn *et al*., 2011). More recently, a study by MacDonald *et al*. indicates that the mono-ubiquitination of the adaptor itself can act as an added anchor point for binding to the Rsp5 ubiquitin ligase (MacDonald *et al*., 2020). We sought to determine if the role of Aly1 and Aly2 in Git1 trafficking was influenced by association with Rsp5 and/or mono-ubiquitination of these α-arrestins. We therefore made use of two classes of mutants for Aly1 and Aly2. The first has each of the ^L^/_P_PxY motifs mutated to ^L^/_P_PxA and is referred to as a ‘PY-less’ mutant. We have shown that these PY-less mutants of the *ALY*s fail to bind Rsp5, are no longer mono-ubiquitinated and fail to execute all clathrin-mediated endocytic functions ascribed to date for these α-arrestins (O’Donnell *et al*., 2013; Prosser *et al*., 2015). The second mutant class has the single lysine needed for α-arrestin mono-ubiquitination mutated to arginine. We mapped this mono-ubiquitination site on Aly1 to lysine 379 and on Aly2 to lysine 392 using mass spectroscopy analyses and demonstrated previously that these K-to-R mutants are no longer ubiquitinated (Hager *et al*., 2018). We next introduced plasmid-borne versions of each of these mutants into *aly1*Δ *aly2*Δ cells and looked to see what impact they had on GPI sensitivity and Git1 trafficking. In keeping with all earlier observations, the ‘PY-less’ mutant versions of either Aly1 or Aly2 were completely non-functional in these assays, and so *aly1*Δ *aly2*Δ cells containing Aly1^PY-less^ or Aly2^PY-less^ remained sensitive to GPI and Git1-GFP was retained at the PM (Figures 6B-F). Interestingly, the Aly1^K379R^ and Aly2^K392R^ mutants that lack their mono-ubiquitination site behaved similarly to wild-type Alys in the GPI growth assays, restoring resistance to GPI, though the Aly2^K392R^ mutant did appear modestly impaired in this assay compared to wild-type Aly2 (Figure 6B). As for Git1 trafficking, the Aly2^K392R^ mutant stimulated trafficking of Git1 to the vacuole under basal conditions as effectively as wild-type Aly2 (Figures 6E-F). However, for Aly1 there was only a very modest increase in trafficking of Git1 to vacuole under these steady-state conditions to begin with and the Aly1^K379R^ mutant was unable to mediate this trafficking (Figure 6C-D). These findings are not so surprising given that Aly1 seems better able to traffic Git1 in response to its ligand GPI and so under these steady-state conditions we see little alteration in trafficking with added Aly1. These data demonstrate that while binding of Aly1 and Aly2 to Rsp5 is needed for their function with Git1, mono-ubiquitination of either of these α-arrestins does not seem to dramatically influence their ability to traffic Git1. It will be interesting in the future to see how the trafficking of assorted cargo may be differentially sensitive to the mono-ubiquitination of the α-arrestin.

**FIGURE 6.**
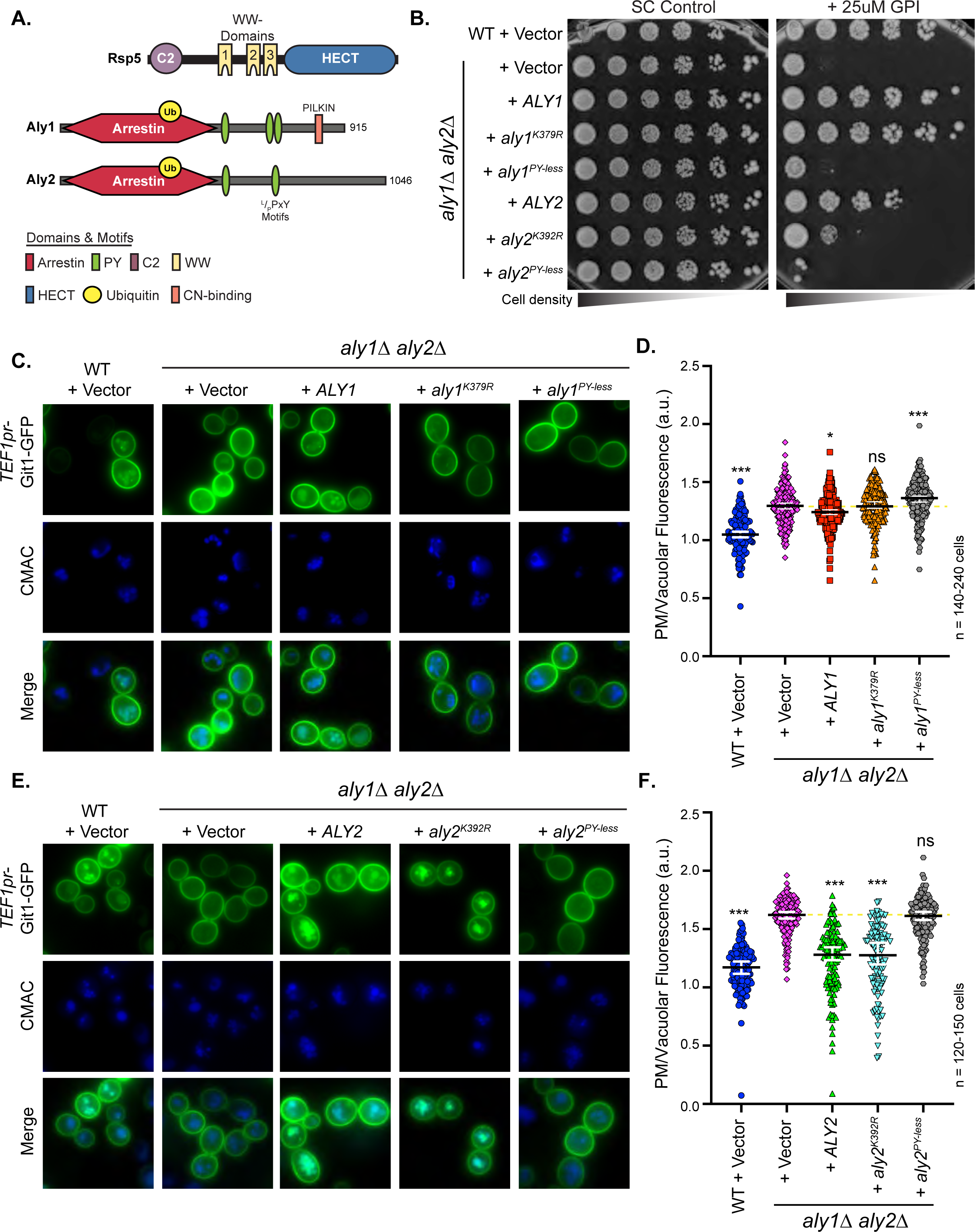
Aly1- and Aly2-mediated trafficking of Git1 requires interaction with Rsp5. *A*, Schematic of Rsp5 and Aly1/Aly2 domain and motif architecture. The site of monoubiquitination on Aly1 and Aly2 is indicated as a yellow circle. *B*, Growth of serial dilutions of WT or *aly1Δ aly2Δ* cells containing a *LEU2*-marked *TEF1pr*-*GIT1*-GFP plasmid and plasmids expressing the α-arrestin variant indicated on SC medium lacking uracil and leucine and containing 25μM GPI is presented. Growth is shown after 2 days of incubation at 30°C. *C and E*, WT or *aly1Δ aly2Δ* cells containing Git1-GFP expressed from the *TEF1* promoter and either a pRS316-derived empty vector expressing nothing or expressing the indicated α-arrestin were imaged by fluorescence microscopy. *D and F*, The PM and vacuolar fluorescence intensities from the cells depicted in C or E, respectively, were quantified using our automated quantification pipeline and the distributions of PM/vacuolar fluorescence ratios in arbitrary units (a.u.) were plotted as scatter plots. The horizontal midline in black represents the median and the 95% confidence interval is represented by the white error bars. Kruskal-Wallis statistical analyses with Dunn’s post hoc test were performed and statistical significance compared to *aly1*Δ *aly2*Δ cells with vector is indicated with asterisks or as not significant (ns) where applicable. In each case, a yellow dashed line is provided that represents the median of the *aly1*Δ *aly2*Δ cells containing a vector control to facilitate comparisons.

### Aly1-mediated trafficking of Git1 is modestly regulated by the protein phosphatase calcineurin

Our earlier studies showed that α-arrestin Aly1 binds the protein phosphatase calcineurin through a canonical 8-amino acid ‘`PxIxIT’` motif, in this case with the sequence `PILKIN` found in the C-terminal tail of Aly1 (Figure 6A) (O’Donnell *et al*., 2013). Calcineurin is a calcium- and calmodulin-dependent protein phosphatase conserved from yeast to man (Roy and Cyert, 2009, 2020). In humans, calcineurin regulates T-cell differentiation, cardiac function and impacts memory and learning (Klee and Haiech, 1980; Klee and Means, 2002). In yeast, calcineurin has an important role in helping cells adapt to cellular stressors by rewiring the transcriptional response via its control of the Crz1 transcription factor’s nuclear translocation (Stathopoulos and Cyert, 1997; Cyert, 2003). However, calcineurin also has an important role in sensing and regulating lipid homeostasis; specifically, by playing a critical role in controlling sphingolipid balance through control of its downstream substrates (Bultynck *et al*., 2006; Tabuchi *et al*., 2006). Given that GPI transport through Git1 can influence PI and sphingolipid levels in cells (Figure 1A), we sought to determine if calcineurin-dependent regulation of Aly1 might influence its role in Git1 trafficking. To do this we made use of mutants that altered CN-mediated regulation of Aly1 in two ways. First, we used Aly1 mutants that lack the CN binding site—either Aly1^ΔPILKIN^, where the site has been deleted, or Aly1^AAAAAA^, where each residue in the PILKIN motif has been mutated to alanine. In each of these mutants, CN can no longer bind to or dephosphorylate Aly1, and therefore Aly1 is retained in its phosphorylated state (O’Donnell *et al*., 2013), providing that the kinases needed to phosphorylate Aly1, which remain undefined, are indeed active. Second, from our earlier studies we identified 5 residues—threonine 250, serine 252, serine 568, serine 569, and serine 573—that showed significant dependence on CN for their regulation based on our mass spectroscopy and mutational analyses (O’Donnell *et al*., 2013). Here we use Aly1 mutants where each of these sites has been converted to an alanine (referred to as Aly1^5A^), to mimic the dephosphorylated state, or a glutamic acid (referred to as Aly1^5E^), to mimic the phosphorylated state (O’Donnell *et al*., 2013). These site mutations bypass the need for kinase activity and directly modulate the CN-regulated sites in question.

Interestingly, cells expressing Aly1 or Aly1^5A^ (mimics constitutively dephosphorylated Aly1) gave rise to the strongest restoration of growth on GPI, while the Aly1^ΔPILKIN^ and Aly1^AAAAAA^ (which cannot be dephosphorylated by CN) gave a slightly more modest rescue of the growth on GPI than wild-type Aly1. Finally, Aly1^5E^ (mimics constitutively phosphorylated form of Aly1) did not rescue growth on GPI at all (Figure 7A). When we examined the localization of Git1 under steady state conditions, we again find that addition of Aly1 to *aly1*Δ *aly2*Δ cells gave only a modest reduction in the PM/vacuole ratio of Git1 that in this experiment was not significantly different from the vector alone (Supplemental Figure 4A-B). The mutant versions of Aly1 that lack the CN-binding site, Aly1^ΔPILKIN^ and Aly1^AAAAAA^, and the CN-regulated phospho-mimetic Aly1^5E^ all failed to significantly alter the PM/vacuole Git1 fluorescence intensities when compared to the vector control in this assay (Supplemental Figure 4A-B). It could be the case that under these conditions the kinase responsible for phosphorylating Aly1 is active and therefore retains these mutants as well as the wild-type Aly1 in an inactive state. However, cells containing Aly1^5A^, which mimics a constitutively dephosphorylated form of Aly1 at these 5 sites, had significantly reduced PM/vacuole ratios compared to *aly1*Δ *aly2*Δ cells with a vector control (Supplemental Figure 4A-B). To better assess Aly1 function under conditions where it would be more active, we next explored GPI-induced trafficking of these same cells and found that when GPI was added Aly1, Aly1^ΔPILKIN^, Aly1^AAAAAA^, Aly1^5A^ and Aly1^5E^ all significantly reduced the PM/vacuole fluorescence ratio in comparison to *aly1*Δ *aly2*Δ containing vector alone (Figure 7B-C). However, the Aly1^5A^ mutation that mimics the dephosphorylated form at these residues, was best able to stimulate vacuolar trafficking of Git1 in these conditions (Figure 7B-C). Taken together, these data further support a model whereby Aly1 plays only a modest role in the steady-state trafficking of Git1, however upon GPI addition Aly1’s role in trafficking Git1 to the vacuole is stimulated. Finally, in response to GPI the most active form of Aly1 for inducing Git1-trafficking to the vacuole is the dephosphorylation mimetic allele at CN-regulated sites, Aly1^5A^. In contrast, the phosphorylated mimetic at these same sites, Aly1^5E^, is unable to rescue growth of *aly1*Δ *aly2*Δ on GPI-containing medium. These results are consistent with CN acting as a modest positive regulator of Aly1-mediated trafficking of Git1.

**FIGURE 7.**
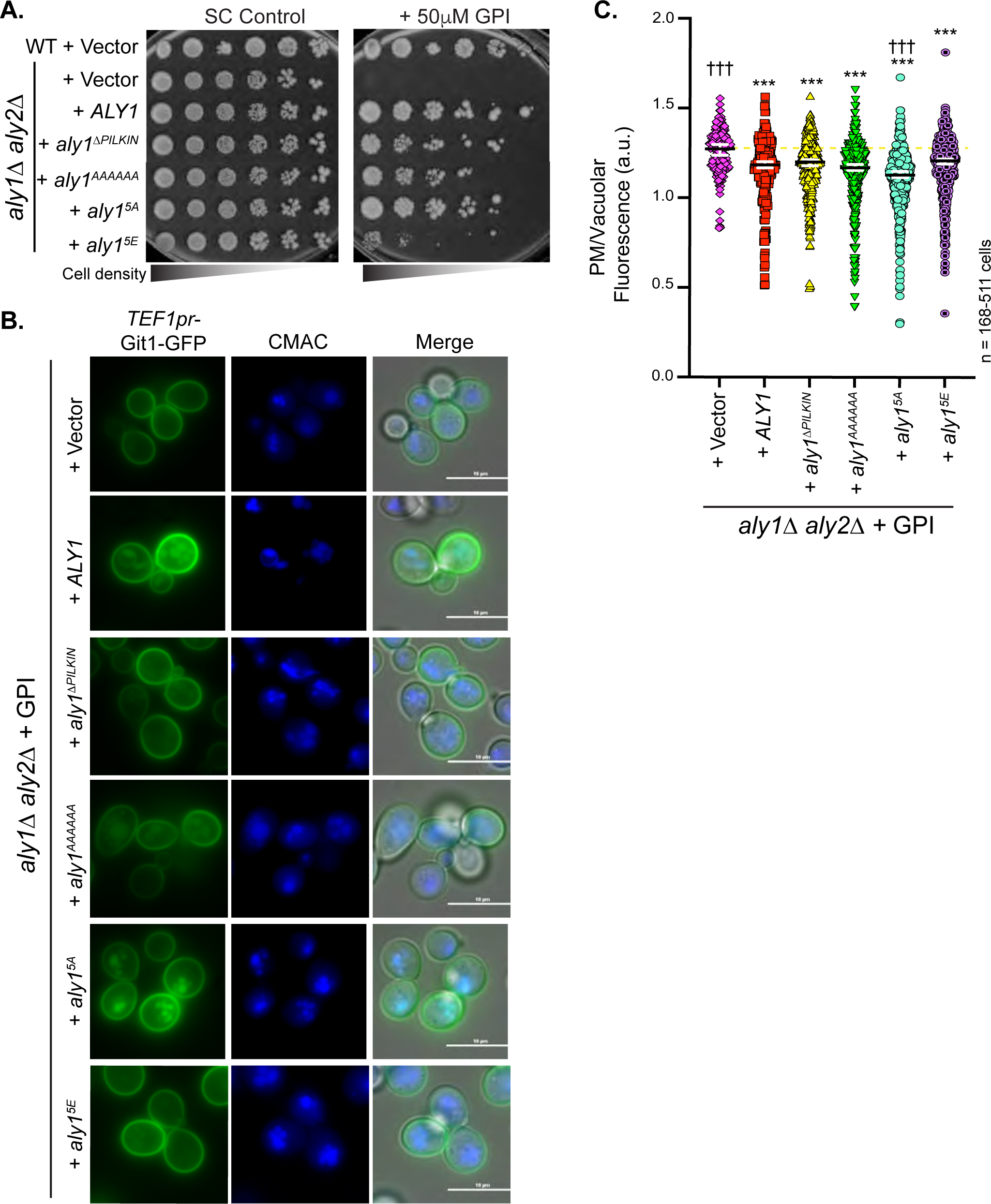
Calcineurin-mediated regulation of Aly1 plays a modest role in Git1 trafficking. *A*, Growth of serial dilutions of WT or *aly1Δ aly2Δ* cells containing a *LEU2*-marked *TEF1pr*-*GIT1*-GFP plasmid and a pRS316 vector expression nothing or expressing the indicated α-arrestins on SC medium lacking uracil and leucine and containing 50μM GPI is shown after 2 days of incubation at 30°C. *B*, Cells lacking ALY1 and ALY2 with a plasmid expressing Git1-GFP from the *TEF1* promoter and a pRS316-empty vector expressing nothing or the indicated α-arrestins were treated with 50μM GPI for 2 h, stained with CMAC and imaged by fluorescence microscopy. *C*, The PM and vacuolar fluorescence intensities from the cells depicted in *B* were quantified using our automated quantification pipeline and the distributions of PM/intracellular fluorescence ratios in arbitrary units (a.u.) were plotted as scatter plots. The horizontal midline in black represents the median and the 95% confidence interval is represented by the white error bars. Kruskal-Wallis statistical analyses with Dunn’s post hoc test were performed and statistical significance compared to the *aly1*Δ *aly2*Δ cells with vector is indicated with asterisks or as not significant (ns) where applicable. The ††† symbols represent p-values of <0.001 and is used to mark comparisons between *aly1*Δ *aly2*Δ expressing *ALY1* and each of the other conditions. In these conditions, most mutants are not statistically different from wild-type Aly1 (no marker shown); Aly1^5A^ is the only mutant that is significantly lower PM/vacuole ratios than wild-type Aly1 while the vector control has significantly higher PM/vacuole ratios. In each case, a yellow dashed line is provided that represents the median of the *aly1*Δ *aly2*Δ cells containing a vector control to facilitate comparisons.

While these data suggest that CN–acting through Aly1–is a modest positive regulator of Git1 trafficking to the vacuole, we were surprised to see that when we deleted the regulatory subunit of CN, Cnb1, Git1 trafficking to the vacuole was greatly increased (Figure 8A-B). In contrast to the modest effects observed when exploring CN-regulation of Aly1, this dramatic change in steady-state localization of Git1 in the absence of CN function suggests that CN plays a largely inhibitory role on the vacuolar sorting of Git1. We sought to define the genetic relationship between the Alys and CN in controlling Git1 localization. When we examined the sensitivity to GPI of *cnb1*Δ cells expressing Git1-GFP, we found that loss of Cnb1 conferred resistance to GPI and trafficking of Git1 to the vacuole was stimulated, an effect that GPI addition did nothing to alter (Figures 8D-E). Importantly, loss of the α-arrestins Aly1 and Aly2 completely blocked the *cnb1*Δ-induced resistance to GPI and trafficking of Git1 to the vacuole (Figure 8D-E), demonstrating that the Aly1/Aly2 regulation of Git1 trafficking is epistatic to that of the Cnb1 regulation of this transporter. We next considered that the function of CN in this regulation may be more robustly influenced by its role in regulating sphingolipid metabolism–which CN does at several points in the sphingolipid biosynthetic pathway (see below)–than controlling α-arrestin-mediated trafficking and describe a more complex interplay between Git1, CN, and protein trafficking as it relates to sphingolipid and phosphoinositide balance.

**FIGURE 8.**
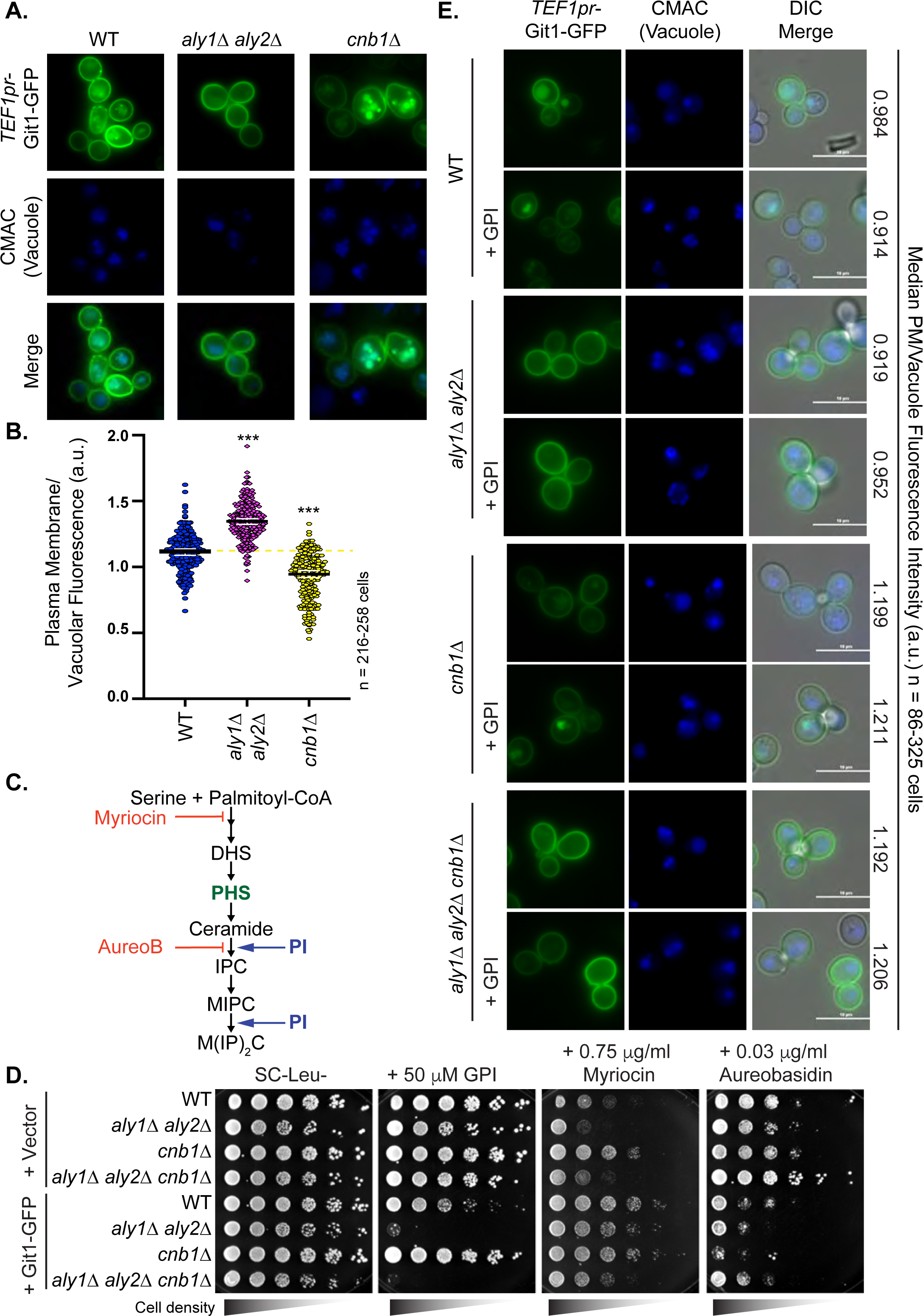
Loss of CN increases Git1 trafficking to the vacuole, but aly1Δ aly2Δ is epistatic to CN regulation. *A*, WT, *aly1Δ aly2Δ*, or *cnb1*Δ cells containing Git1-GFP expressed from the *TEF1* promoter were stained with CMAC and imaged by fluorescence microscopy. *B*, The PM and vacuolar fluorescence intensities from the cells depicted in A were quantified using our automated quantification pipeline and the distributions of PM/vacuolar fluorescence ratios in arbitrary units (a.u.) were plotted as scatter plots. The horizontal midline in black represents the median and the 95% confidence interval is represented by the white error bars. Kruskal-Wallis statistical analyses with Dunn’s post hoc test were performed and statistical significance compared to the wild-type cells is indicated with asterisks. A yellow dashed line is provided that represents the median of wild-type cells to facilitate comparisons. *C,* Schematic of the sphingolipid biosynthetic pathway demonstrating the points where myriocin and aureobasidin (AureoB) impair the pathway (red lines) and the sites where PI is incorporated into the complex base production in the lower half of the pathway (PI with blue arrows). The intermediates in the pathway are shown but the enzymes are not indicated. Abbreviations in this diagram are as follows: DHS (dihyrdosphingosine), PHS (phytosphingosine), IPC (inositolphosphoryl-ceramide), MIPIC (mannosylinositolphosphoryl-ceramide), and M(IP)_2_C (mannosyl-di-inositolphosphoryl ceramide). For simplicity, some intermediates are not shown and instead, multiple arrowheads are provided for those steps where two or more enzymatic steps have been condensed. *D*, Growth of serial dilutions of the cells indicated containing either a vector or a *LEU2*-marked *TEF1pr*-*GIT1*-GFP plasmid on SC medium lacking leucine and containing the indicated concentrations of GPI, myriocin, or aureobasidin are shown after 2-4 days of incubation at 30°C. *E*, WT, *aly1Δ aly2Δ*, *cnb1*Δ, or *aly1Δ aly2Δ cnb1*Δ cells containing Git1-GFP expressed from the *TEF1* promoter were treated with 50μM GPI for 2h, stained with CMAC and imaged by fluorescence microscopy. Automated quantification of the PM and vacuole fluorescence intensities was performed and the median PM/vacuole fluorescence intensity for each is presented in arbitrary units (a.u.) on the right side of the figure.

### Aly-dependent alterations in sphingolipid and phosphoinositide metabolism

The GPI internalized via Git1 releases free inositol that is incorporated into PI. Since PI is a precursor to sphingolipid and phosphoinositide production (Figure 1A and Figure 8C), we probed each of these pathways for possible alterations in function when either α-arrestin-mediated or general protein trafficking were perturbed and also queried regulation of this pathway by CN. First, we examined the sensitivity of cells lacking Aly1 and/or Aly2 to myriocin, a drug that inhibits the enzymes Lcb1 and Lcb2 which act in the first step of the *de novo* sphingolipid synthesis pathway (Figure 8C), and aureobasidin, a drug that blocks the phosphatidylinositol:ceramide phosphoinositol transferase Aur1’s ability to convert ceramide to inositol phosphorylceramide (IPC) in sphingolipid synthesis (Figure 8C) (Angus and Lester, 1972; Dickson *et al*., 2006; Epstein *et al*., 2012; Klug and Daum, 2014; Montefusco *et al*., 2014). Since elevated ^3^H-GPI uptake leads to higher conversion of the inositol label into PI (Figure 10C) and PI is required for complex sphingolipid biosynthesis, we hypothesized that loss of Aly1 and Aly2 might make cells more resistant to inhibition of sphingolipid synthesis. However, this is not what we observed. Loss of Aly1 and Aly2 resulted in modest sensitivity to myriocin and aureobasidin (Figure 8D), suggesting that this mutant may be defective in maintaining sphingolipid balance via mechanisms that go beyond Git1 trafficking. The myriocin sensitivity of *aly1*Δ *aly2*Δ cells was rescued by addition of phytosphingosine (PHS) (Supplemental Figure 5A), consistent with defective flux through this pathway (Figure 8C). Interestingly, other endocytic mutants, including *vrp1*Δ and *end3*Δ, or ESCRT mutants, like *vps4*Δ and *vsp27*Δ, which would each ultimately increase abundance of membrane proteins at the cell surface either by blocking endocytosis or by preventing their degradation in the MVB pathway and allowing ample opportunity for recycling, similarly showed sensitivity to myriocin that can, in all but *end3*Δ cells, be partially rescued by PHS addition (Supplemental Figure 5A). Thus, defective protein trafficking may lead to impaired sphingolipid homeostasis. Given the importance of sphingolipids in the plasma membrane and other intracellular membranes, it is not surprising that defects in this membrane’s turnover could change sphingolipid balance. Indeed, many trafficking mutants have been associated with defective sphingolipid balance, including the Golgi-associated retrograde protein (GARP) complex (Frohlich *et al*., 2015), regulators of Niemann-Pick type C disease (Newton *et al*., 2018), and Rab proteins (Choudhury *et al*., 2002). Based on these data it appears that the α-arrestins Aly1 and Aly2 influence sphingolipid homeostasis as well, however the linkage to Git1 regulation, as opposed to other effects, is unclear.

We next examined *cnb1*Δ cells and found that they were sensitive to myriocin and resistant to aureobasidin (Figure 8D), consistent with a role for CN as an effector of the sphingolipid biosynthesis pathway (Dunn *et al*., 1998; Bultynck *et al*., 2006; Tabuchi *et al*., 2006; Muir *et al*., 2014). However, the *aly1*Δ *aly2*Δ *cnb1*Δ cells were wild-type like in their sensitivity to myriocin but were completely resistant to aureobasidin, suggesting a more complex interplay between these α-arrestins and CN than previously described. Given that CN impinges upon the sphingolipid biosynthesis pathway at multiple points, including i) dephosphorylation of Lac1/Lag1, the ceramide synthases involved in generating ceramide from long chain bases in sphingolipid biosynthesis, ii) regulation of Csg2, a calcium sensitive gene and one of two paralogous mannosylinositol phosphorylceramide (MIPC) synthase catalytic subunits, and iii) dephosphorylation of the Slm proteins, which are PI(4,5)P_2_ binding proteins that operate downstream of the phosphatidylinositol-4-phosphate 5-kinase Mss4 in controlling actin cytoskeleton dynamics, TORC2 activity and sphingolipid metabolism (Dunn *et al*., 1998; Bultynck *et al*., 2006; Tabuchi *et al*., 2006; Muir *et al*., 2014), it is to be expected that the entire relationship here will not be solely explained via the α-arrestins. Considering these broad roles in sphingolipid regulation for CN, it is very interesting that the *aly1*Δ *aly2*Δ completely reverses the aureobasidin sensitivity for *cnb1*Δ cells (Figure 8D). To begin to determine if any of these phenotypes were linked to Git1 trafficking, we over-expressed Git1-GFP in each of these mutants and found that over-expression of our Git1-GFP construct increased resistance to myriocin in all but the *cnb1*Δ cells, while increasing sensitivity to aureobasidin (Figure 8D). Again, these findings are not what one would expect since elevated Git1 should increase GPI uptake and elevate PI levels, which is known to promote sphingolipid biosynthesis. Thus, while these data further define a role for Git1, CN and endocytic trafficking in influencing sphingolipid balance, the details of the interplay between these factors and the underlying mechanism for these altered sensitivities to sphingolipid inhibition remain to be fully elucidated.

In addition to feeding PI into the sphingolipid biosynthetic pathway, uptake of GPI through Git1 and its subsequent release of inositol could elevate PI to cause changes in membrane phospholipid production. To survey the phospholipid distributions in cells where Aly1 and Aly2 were lost, we examined the localization and abundance of probes for phosphatidylserine (PS, Lact-C2-GFP probe) as well as phosphatidylinositol 4-phosphate (PI(4)P, GFP-2xPH-Osh2 and GFP-P4Mx1 probes), phosphatidylinositol 3-phosphate (PI(3)P, GFP-FYVE-Eea1 probe), and phosphatidylinositol 4,5-bisphosphate (PI(4,5)P_2_, GFP-2xPH-Plc*δ*). We found no significant changes in PI(4)P or PI(4,5)P_2_ in the absence of α-arrestins Aly1 or Aly2 (Supplemental Figure 6A-C). However, there were dramatic changes in PI(3)P and PS in the *aly1*Δ *aly2*Δ mutants compared to wild-type cells (Figure 9A-D). Interestingly, the PI(3)P, which typically marks the endosomal compartment in wild-type cells, was significantly brighter and distributed to the limiting membrane of the vacuole in the absence of the Alys. In addition, the PS at the cell surface was much higher in these mutants (Figure 9A-C). Though it is somewhat unclear how these specific phospholipid species, and not others, would be influenced by α-arrestins, we sought to determine if alteration of PI(3)P could in any way be linked to GPI uptake through Git1. We treated cells with GPI and monitored GFP-FYVE-Eea1 localization and abundance changes. We found that even in wild-type cells treated with GPI, PI(3)P-associated fluorescence increased with brighter PI(3)P signal on the limiting membrane of the vacuole (Figure 10A-B). In the absence of *ALY1* and *ALY2*, not only did cells start out with more PI(3)P on the vacuole membrane, the abundance further increased upon GPI addition (Figure 10A-B). To further assess the linkage between GPI uptake and PI conversion as possibly fueling these changes in phospholipid balance, we provided cells with [^3^H]inositol-GPI and monitored the presence of the inositol label across extracellular, soluble intracellular, and membrane fractions (Table 1). These assays were performed using the endogenous Git1 transporter under growth conditions that would allow for *GIT1* expression (see Methods). As shown in Table 1 and Figure 10C, *aly1Δ aly2Δ* cells incorporated a greater percentage of inositol label from exogenous GPI into the cell as both a TCA-extractable (water-soluble) fraction and into a particulate membrane fraction as compared to wild-type cells during the labeling period. Roughly 2.5 times the amount of soluble intracellular inositol label and 4 times as much PI was found in *aly1Δ aly2Δ* as compared to wild-type (Table 1). This result is consistent with increased stability, and hence activity, of Git1 at the PM in *aly1Δ aly2Δ* cells. It is important to note that the labeled metabolites detected, including PI, do not represent total internal pools, but rather represent only those molecules that received inositol label from exogenous [^3^H]-GPI. In addition to this pool, *de novo* synthesis of inositol will be a major source of unlabeled inositol and that pool is not monitored by these assays (Henry *et al*., 2012; Henry *et al*., 2014). These data confirm the known linkage between GPI uptake and incorporation of released inositol into PI and show that the α-arrestins Aly1 and Aly2 play an important role in this regulation (Table 1) (Patton *et al*., 1995; Patton-Vogt and Henry, 1998; Almaguer *et al*., 2003; Almaguer *et al*., 2004). This is not a completely unexpected functional link between GPI uptake, its conversion to PI and an increase in PI(3)P, as these pathways are well-documented. However, our findings do not demonstrate, nor do we mean to suggest, that all of the aberrant PI(3)P accumulation on vacuoles *aly1*Δ *aly2*Δ cells is due to GPI uptake via Git1. In fact, under the conditions used for the PI(3)P imaging assays we would expect very little Git1 to be at the cell surface and so there must be additional mechanisms by which *aly1*Δ *aly2*Δ causes elevated PI(3)P that remains to be explored.

**FIGURE 9.**
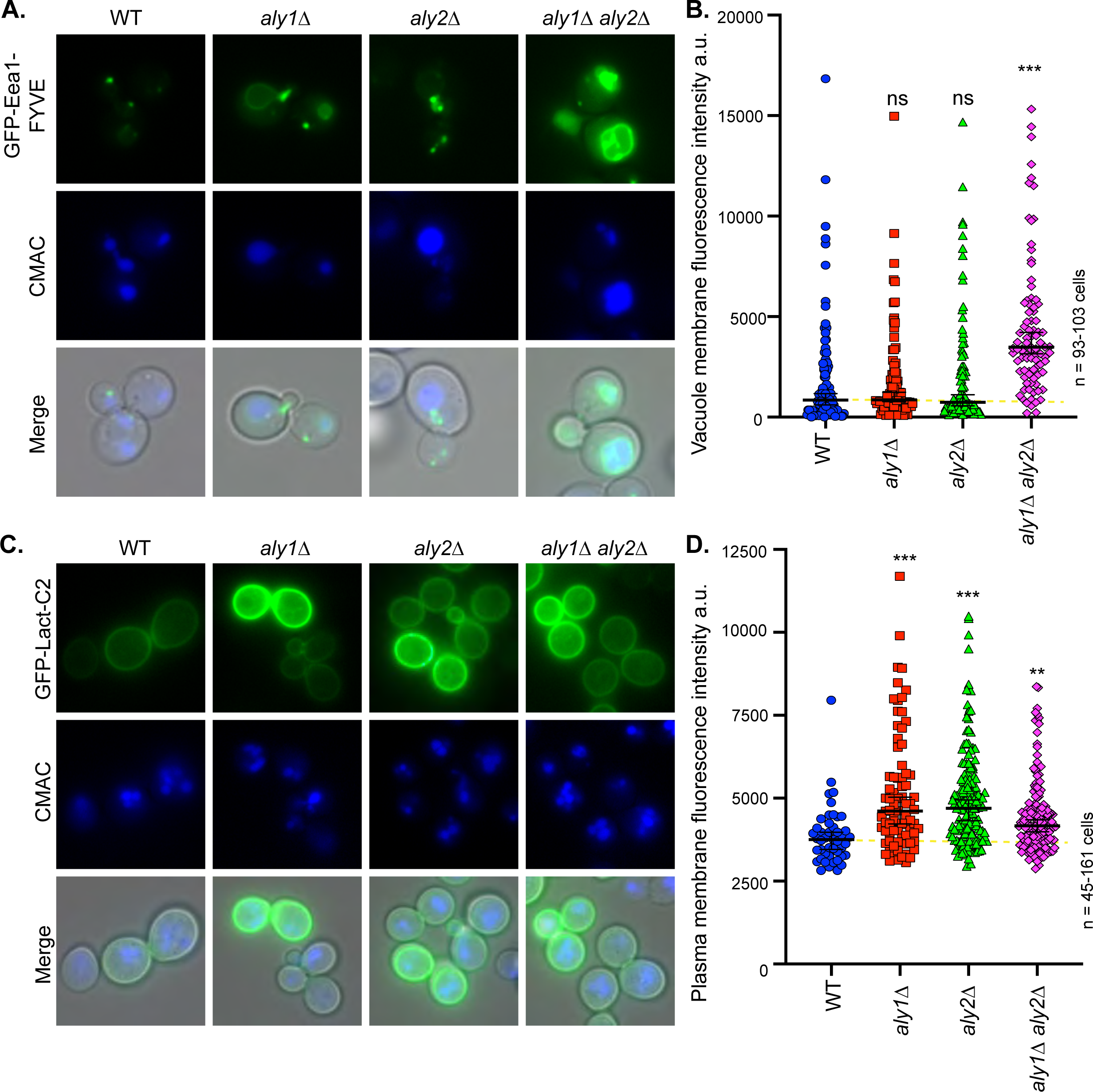
Loss of Aly1 and Aly2 disrupts phospholipid balance in the endomembrane system. *A*, The indicated cells were transformed with the plasmid expressing a GFP-fused FYVE domain of Eea1, which marks PI(3)P. Cells were stained with CMAC and imaged by fluorescence microscopy. *B*, The GFP fluorescence at the limiting membrane of the vacuole from the images in A was quantified by manually drawing a line around the outside of the CMAC stained vacuoles and overlaying this mask onto the GFP channel. The quantified fluorescence intensities (arbitrary units, a.u.) are presented as scatter plots. The horizontal midline in black represents the median and the 95% confidence interval is represented by the black error bars. Kruskal-Wallis statistical analyses with Dunn’s post hoc test were performed and statistical significance compared to the wild-type cells is indicated with asterisks or as not significant (ns) where applicable. A yellow dashed line is provided that represents the median of wild-type cells to facilitate comparisons. *C*, The indicated cells were transformed with the plasmid expressing a GFP-fused C2 domain of the Lact protein, which marks PS. Cells were stained with CMAC and imaged by fluorescence microscopy. *D*, The GFP fluorescence at the plasma membrane from the images in C was quantified using our automated pipeline. The quantified fluorescence intensities (arbitrary units, a.u.) are presented as scatter plots. The horizontal midline in black represents the median and the 95% confidence interval is represented by the black error bars. Kruskal-Wallis statistical analyses with Dunn’s post hoc test were performed and statistical significance compared to the wild-type cells is indicated with asterisks or as not significant (ns) where applicable. A yellow dashed line is provided that represents the median of wild-type cells to facilitate comparisons.

**Table 1.**
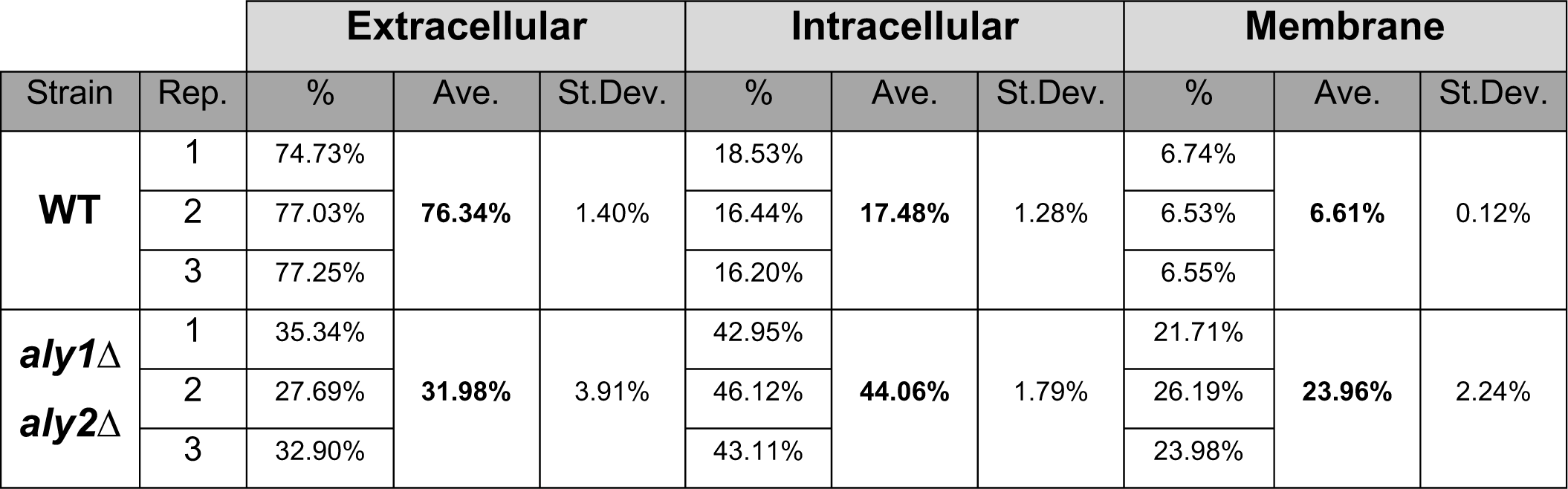
Uptake and conversion of [^3^H]-GPI into [^3^H]-PI. Strains were grown in the presence of 5μM [^3^H-inositol]-GPI (see Methods) and then extracellular, intracellular and membrane fractions were isolated. Lipids were extracted from the membrane fraction and resolved by TLC as described in *Methods*. The average (Ave.) and standard deviation (St.Dev.) of three individual cultures (biological triplicates) are presented. It should be noted that this experiment was repeated with label of lower specific activity and similar results were obtained.

## Discussion

Our findings demonstrate that α-arrestins Aly1 and Aly2 regulate the endocytosis of the Git1 transporter, may contribute to its intracellular sorting, and ultimately stimulate Git1 trafficking to the vacuole. We show for the first time that Git1 is internalized in response to the presence of its substrate, GPI, and both basal and ligand-induced internalization is dependent upon the Alys. However, there appears to be an interesting division of labor between Aly1 and Aly2, with Aly2 doing the lion’s share of the trafficking under steady-state conditions, while the role of Aly1 in trafficking Git1 appears to be more selectively stimulated in response to GPI. Consistent with earlier studies of the Git1 transporter (Almaguer *et al*., 2006), its internalization depends upon clathrin as we can block Git1 endocytosis using mutations in early clathrin-coated pit assembly, by preventing actin polymerization using LatA or with mutants lacking the clathrin adaptor complex AP-2. This latter finding is particularly interesting since there are very few endocytic cargos associated with AP-2 in yeast and we have previously found that Aly1 and Aly2 can each interact directly with AP-2 (O’Donnell *et al*., 2010). Perhaps the Alys contribute together with AP-2 to regulate the endocytosis of Git1, which would be an intriguing point for further exploration. Finally, both Aly1 and Aly2 require their interaction with the ubiquitin ligase Rsp5 to control Git1 trafficking, though neither one’s function appears to be dramatically hampered by loss of their respective mono-ubiquitination sites.

While endocytosis plays a clear role in trafficking Git1 to the vacuole, there may be contributions via intracellular sorting as well. Evidence in support of an intracellular sorting mechanism driving Git1 to the vacuole includes the fact that the *end3*Δ or LatA treated cells do not retain as much Git1 at the PM as *aly1*Δ *aly2*Δ cells, suggesting that these α-arrestins may help with sorting Git1 to the vacuole in addition to regulating endocytosis. In particular, LatA is effective at blocking the impact of Aly2 on Git1 localization but does not seem as effective at blocking Aly1-mediated trafficking of Git1 to the vacuole. This suggests a functional dichotomy between these two α-arrestins. In addition, the block to vacuolar trafficking of Git1 in *aly1*Δ *aly2*Δ upon GPI addition is not complete, suggesting that other adaptors or pathways are also important. Indeed, we find that there is a role for the Buls in regulating Git1 trafficking, specifically of Bul2 in altering cell sensitivity to GPI and this is not the first example of an overlap in function between the Alys and the Buls (Crapeau *et al*., 2014). Loss of Bul2 increases cell sensitivity to GPI, and yet surprisingly, we find that loss of Buls diminishes Git1 abundance, reducing the cell surface levels of Git1 suggesting that the Buls are responsible for altering Git1 activity in some other way than simply allowing for its retention at the PM. We further provide evidence that additional α-arrestins may play minor roles since i) *aly1*Δ *aly2*Δ is not as effective as 9ArrΔ cells (where the Buls are intact) in retaining endogenous Git1 at the cell surface and ii) in the 9ArrΔ cells, Aly1 is more effective at trafficking Git1 under basal conditions than it is in the *aly1*Δ *aly2*Δ cells, suggesting that it may still be competing with other α-arrestins for access to the cargo in this latter cell type.

Earlier studies have shown that α-arrestin Aly1 is dephosphorylated by the protein phosphatase CN and that this activity is needed for optimal Aly1-mediated endocytosis of the aspartic acid permease Dip5 in response to excess ligand (O’Donnell *et al*., 2013). However, the role of CN-regulation in Git1 trafficking is more complex than this simple model, perhaps due to CN’s broader functions in regulating sphingolipid homeostasis. Somewhat surprisingly, mutants of Aly1 that lack the CN-binding site had only a modest impact on Aly1-mediated steady-state or GPI-induced trafficking of Git1, perhaps indicating that the as-yet unknown kinase responsible for countering CN function is not fully active under these conditions. However, a mutant of Aly1 that side-steps the need for kinase function, like the one where the CN-regulated phosphorylation sites were all mutated to alanines to mimic a constitutively dephosphorylated state, significantly improved Git1 trafficking to the vacuole in both steady-state and ligand-induced conditions. This finding is in keeping with the idea that dephosphorylation—or in this case conversion of phosphorylated residues to ones that cannot be modified— improves an α-arrestin’s ability to induce trafficking to the vacuole. It was surprising then to see that loss of the regulatory subunit of CN resulted in robust steady state trafficking of Git1 to the vacuole, indicating that CN has an inhibitory role in Git1 trafficking. Further, loss of the α-arrestins Aly1 and Aly2 in the *cnb1*Δ strains completely blocked this trafficking, demonstrating that the negative regulation CN exerts on this pathway occurs, either directly or indirectly, via the α-arrestins. Perhaps the altered sensitivities to myriocin and aureobasidin, two drugs that inhibit steps in the sphingolipid biosynthetic pathway, upon Git1 over-expression is linked to this element of CN regulation. Indeed, complex sphingolipid production is reduced under conditions where cells are starved for inositol and/or PI and elevated in conditions where there is excess inositol and/or PI (Henry *et al*., 2014). When Git1 is retained at the cell surface it should provide added GPI and, subsequent to its breakdown, inositol for PI synthesis. This in turn should stimulate sphingolipid production and give resistance to drugs that impede the pathway (Henry *et al*., 2012; Henry *et al*., 2014). However, counter to the anticipated model we find increased sensitivity to aureobasidin when Git1 is over-expressed in cells. The reasoning for this phenotype is unclear, but it is important to note that these inhibitor studies were performed in the absence of exogenous GPI (Figure 12D) in order to avoid its toxicity phenotype. In the absence of exogenous supplementation, the extracellular levels of GPI available for transport are estimated to be 1-2 μM per OD in a wild type cell (Angus and Lester, 1972; Patton-Vogt and Henry, 1998) and therefore may not have a dramatic impact on PI levels. Instead, the observed sensitivity result may be related to a broader role for Aly1 and Aly2 in altering trafficking of multiple membrane proteins and/or changing the intracellular signaling landscape. In the end, while our data suggest interesting connections between α-arrestins, Git1, CN and sphingolipid balance, the nature of the mechanistic links remains to be resolved.

A striking finding from this work is the dramatic alteration in certain phospholipid species associated with loss of Alys and exacerbated by addition of GPI as monitored through imaging. While PI(3)P is typically sequestered to endosomal membranes, we show a substantial increase in PI(3)P on the limiting membrane of the vacuoles in *aly1*Δ *aly2*Δ cells. While this phenotype can be linked at least in part to the activity of Git1, since addition of GPI to either wild-type or *aly1*Δ *aly2*Δ cells increased PI(3)P on vacuolar membranes, it seems unlikely that all of the PI(3)P accumulation in the absence of Alys is due solely to Git1. In yeast cells, PI conversion to PI(3)P occurs through the action of Vps34, the phosphatidylinositol 3-kinase localized largely to the vacuole membrane (Schu *et al*., 1993). Perhaps this enrichment in PI(3)P in *aly1*Δ *aly2*Δ cells, which is exacerbated upon GPI addition, is due to hyperactivity of Vps34. Conversely, the Fig4 protein, which is the PI(3,5)P_2_ phosphatase localized to vacuole and nuclear periphery (Gary *et al*., 2002), is responsible for converting PI(3,5)P_2_ back into PI(3)P. Again, hyperactivity of this enzyme could similarly contribute to the elevated PI(3)P, though given the relatively small pool of PI(3,5)P_2_ in cells this seems somewhat less likely (Hasegawa *et al*., 2017).

Conversion of PI(3)P back into PI is regulated by the PI(3)P phosphatase, Ymr1, and the synaptojanin-like phosphatases, Sjl2 and Sjl3 (Parrish *et al*., 2004). In mutants lacking Ymr1 and Sjls there is aberrant accumulation of PI(3)P on the limiting membrane of the vacuole and an associated vacuole fragmentation phenotype (Parrish *et al*., 2004). While there is no readily observable vacuole fragmentation effect in *aly1*Δ *aly2*Δ, a loss of Ymr1/Sjl activity could also contribute to the elevated PI(3)P on the vacuole surface we are observing. It will be interesting to see what influence GPI has on the activity of these many PIP regulatory pathways, as it seems feasible that the mechanism by which Git1 retention at the cell surface leads to GPI-induced toxicity could be through alteration of these pathways. Indeed, maintaining the balance of PIPs in cells is critical as defects in phospholipid metabolism and regulation can have toxic consequences for the cell (Strahl and Thorner, 2007; Shyu *et al*., 2019).

Though we know from our studies that PI(3)P is elevated, typically PI(3)P is converted to PI(3,5)P2 on the limiting membrane of the vacuole, thus raising the question of how this shift in PI(3)P at this location impacts PI(3,5)P_2_. It is difficult to monitor PI(3,5)P_2_ without specialized radiolabeling assays; however, loss of PI(3,5)P_2_, as results when Fab1, the vacuole localized PI(3)P kinase that generates PI(3,5)P_2_, or Vac7, an integral vacuolar membrane protein that activates the Fab1 kinase, are mutated is associated with an enlarged vacuole morphology (Gary *et al*., 2002). Since the vacuoles in *aly1*Δ *aly2*Δ cells do not display this enlarged morphology, it seems unlikely that loss of Alys is leading to dramatic reductions in PI(3,5)P_2_.

While there are distinct changes in PI(3)P observed, it is interesting that we do not see any significant elevation in PI(4)P or PI(4,5)P_2_ which can be produced from PI via the activity of PI 4-kinases Pik1 or Stt4 followed by the PI(4)P 5-kinase Mss4, respectively (Strahl and Thorner, 2007). Perhaps the fact that these PIPs are not disrupted suggests a selective shunting of the PI from GPI into PI(3)P, reflecting a specific defect in the *aly1*Δ *aly2*Δ cells, or suggesting a more rigorous feedback mechanism to maintain PI(4)P and PI(4,5)P_2_ levels within the Golgi and at the PM than is operational for PI(3)P in endosomes. Also, small changes in the levels of these minor lipids could be missed by a purely imaging approach.

It will be interesting in future studies to determine if loss of other α-arrestins contributes to shifts in phospholipid balances inside of cells and to uncover the underlying mechanism behind this phospholipid imbalance. While many components that disrupt protein trafficking alter phospholipid ratios in the cell, the link here between excess GPI uptake due to Git1 retention at the cell surface, conversion of internalized GPI to PI and specific enrichment in PI(3)P at the vacuolar membrane in the absence of α-arrestins Aly1 and Aly2 represent significant new findings which demonstrate that α-arrestins contribute to phospholipid balance in a previously unanticipated way.

## Supporting information

Robinson et al Supplemental

## Acknowledgements

This research was funded by the National Sciences Foundation (MCB CAREER 1902859 and 1553143 to A.F.O.) and the National Institutes of Health (R15 GM 104876 to J.P.V.). The work was also supported by start-up funds from the Depts of Biological Sciences at Duquesne University and the Univ. of Pittsburgh to A.F.O. We thank members of the O’Donnell lab, especially Nejla Ozbaki-Yagan for strain construction, and the Patton-Vogt, as well as the Pittsburgh Area Yeast community, for their critical feedback on this work. We thank Dr. Gerry Hammond (Univ. of Pittsburgh) for the PI(4,5)P_2_ probe.

## Materials and Methods

### Yeast strains and growth conditions

All yeast strains used in this study are derived from the BY4741 genetic background of *S. cerevisiae* (S288C in origin) and are described in detail in Table 2. Yeast were grown in synthetic complete (SC) medium lacking the appropriate nutrient for plasmid maintenance as described in (Johnston *et al*., 1977) or YPD medium where indicated. Liquid medium was filter sterilized and solid medium for agar plates had 2% agar w/v added before autoclaving. Yeast cells were grown at 30°C unless otherwise noted.

**Table 2.**
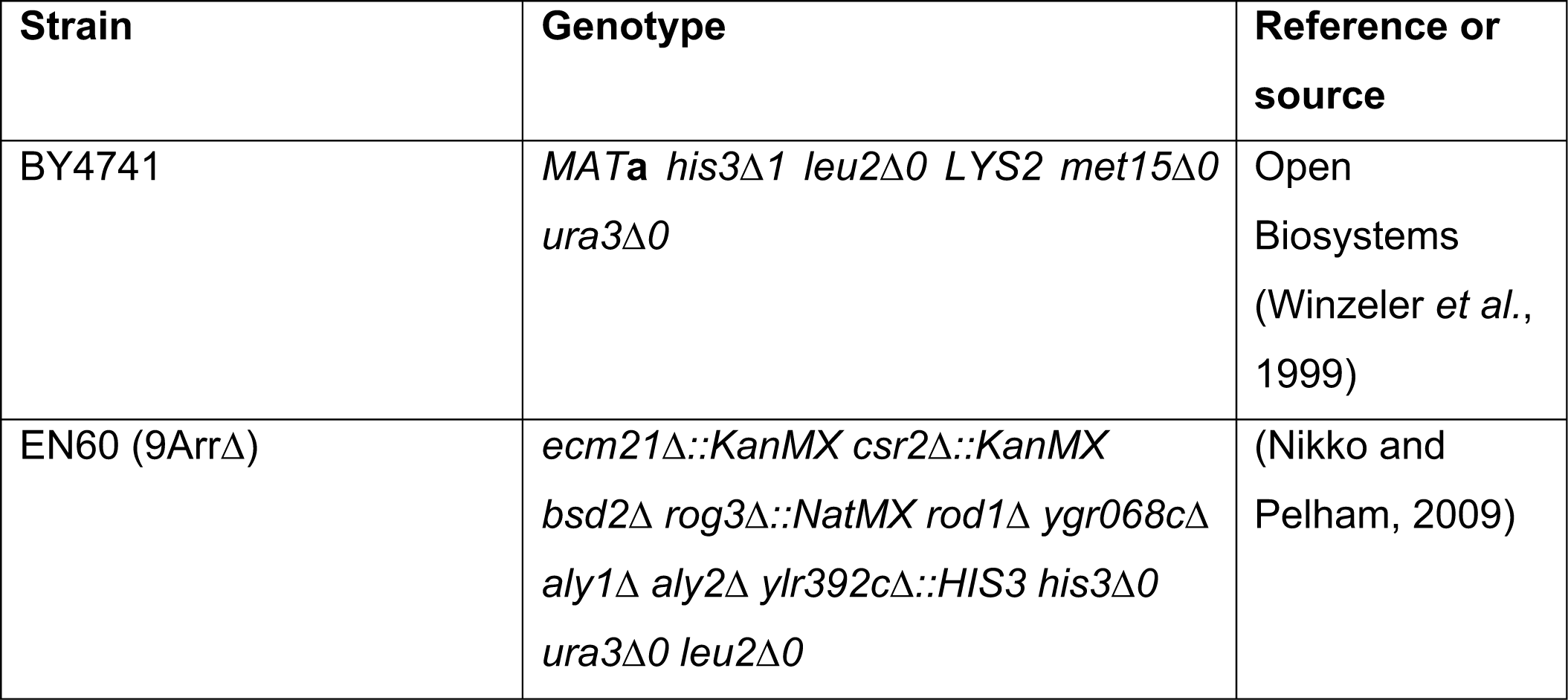
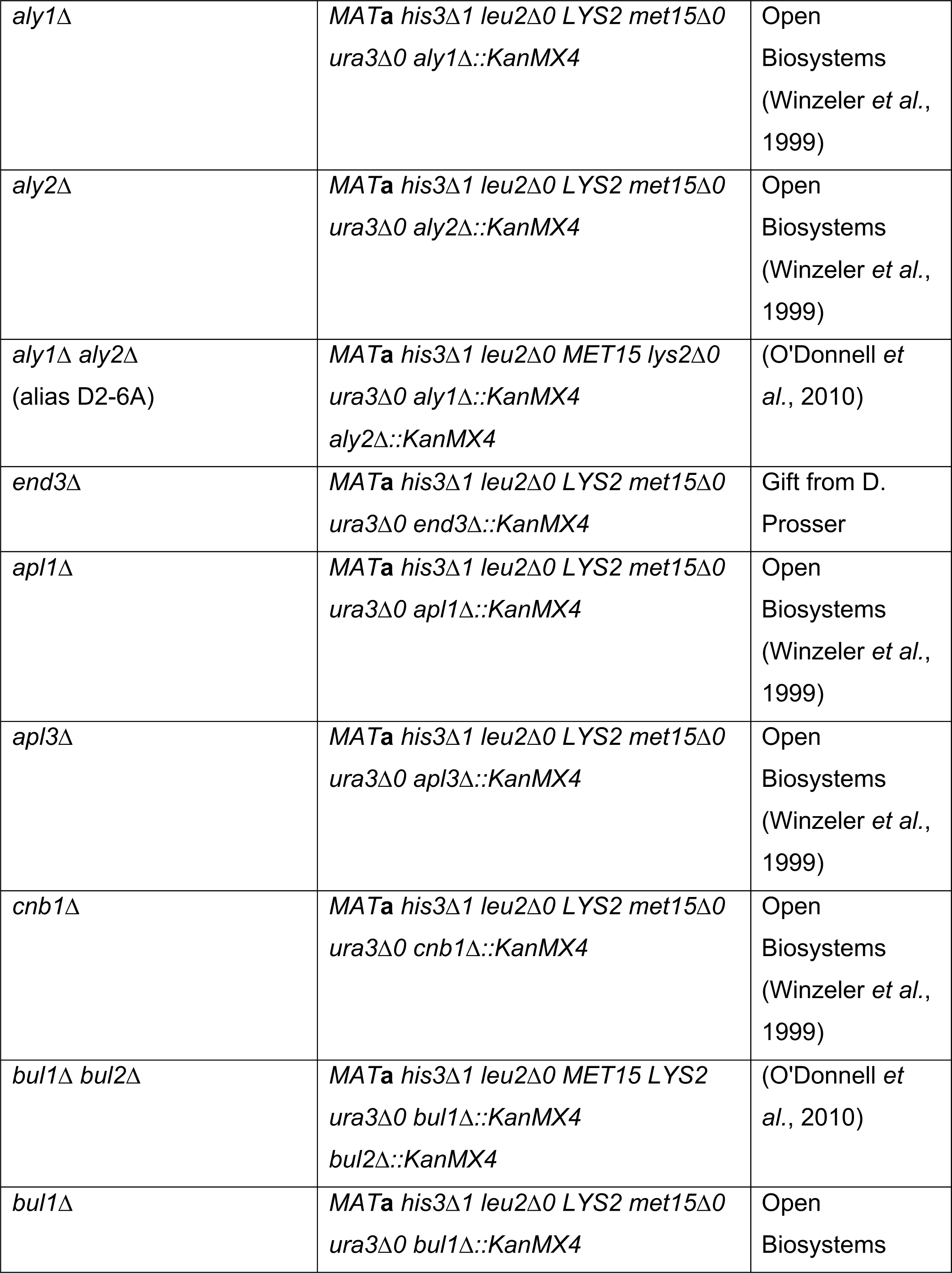
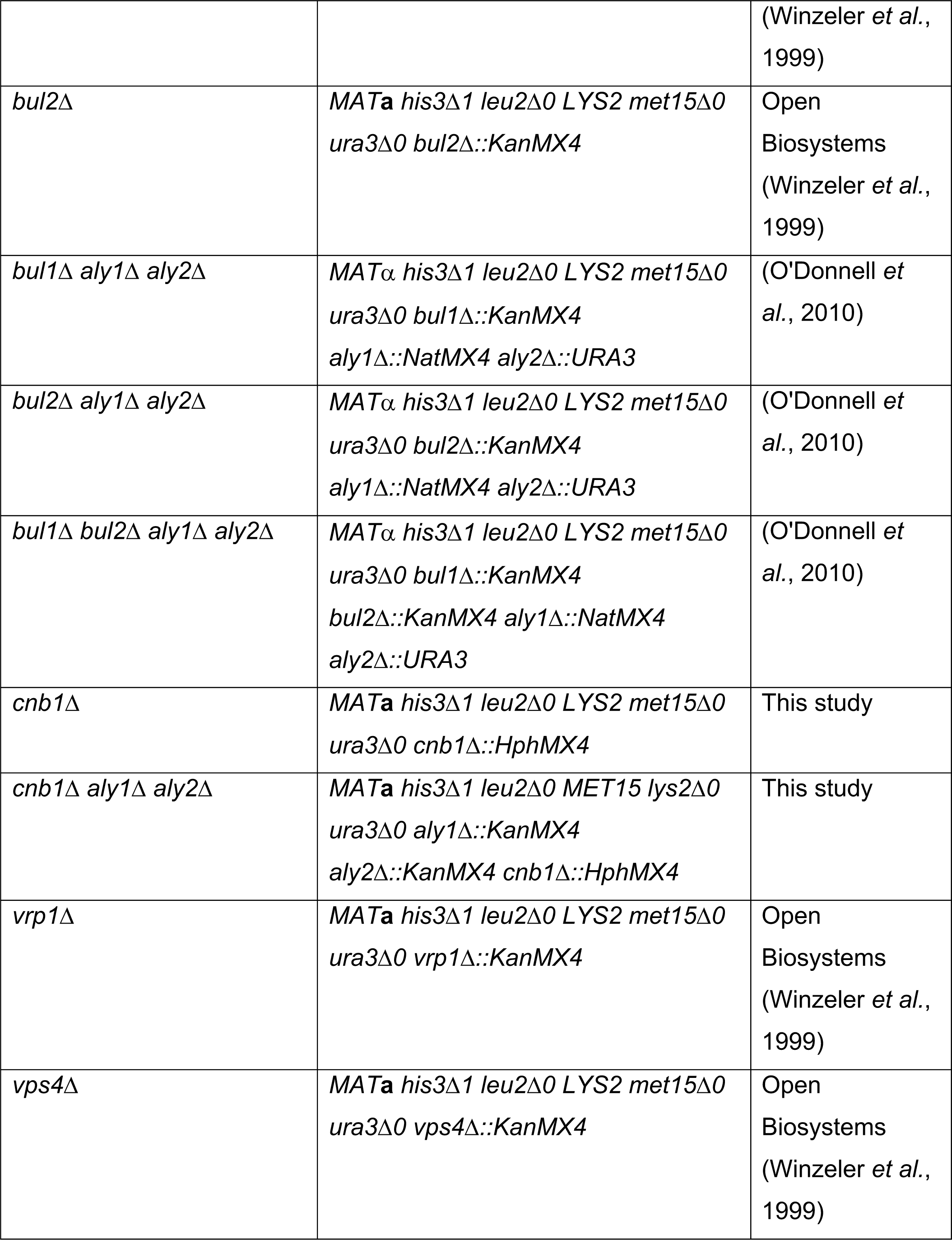
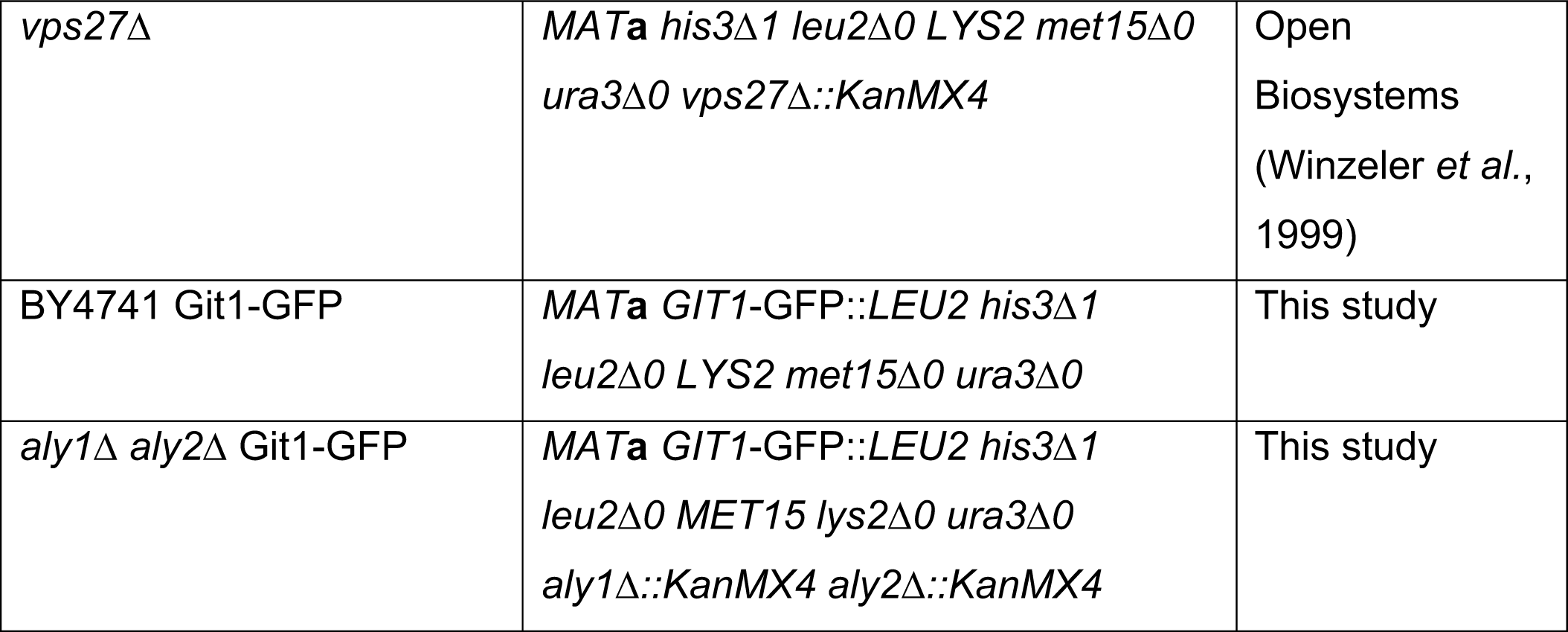
Yeast Strains.

### Plasmid and DNA manipulations

Plasmids used in this work are described in Table 3. PCR amplifications for generating plasmid constructs were performed using Phusion High Fidelity DNA Polymerase (ThermoFisher Scientific, Waltham, MA) and sequence insertions were verified using Sanger sequencing (Genewiz, South Plainfield, NJ). Plasmids were transformed into yeast cells using the lithium acetate method (Gietz and Schiestl, 2007)and selected for on the appropriate SC medium lacking the specific nutrient marker.

**Table 3.**
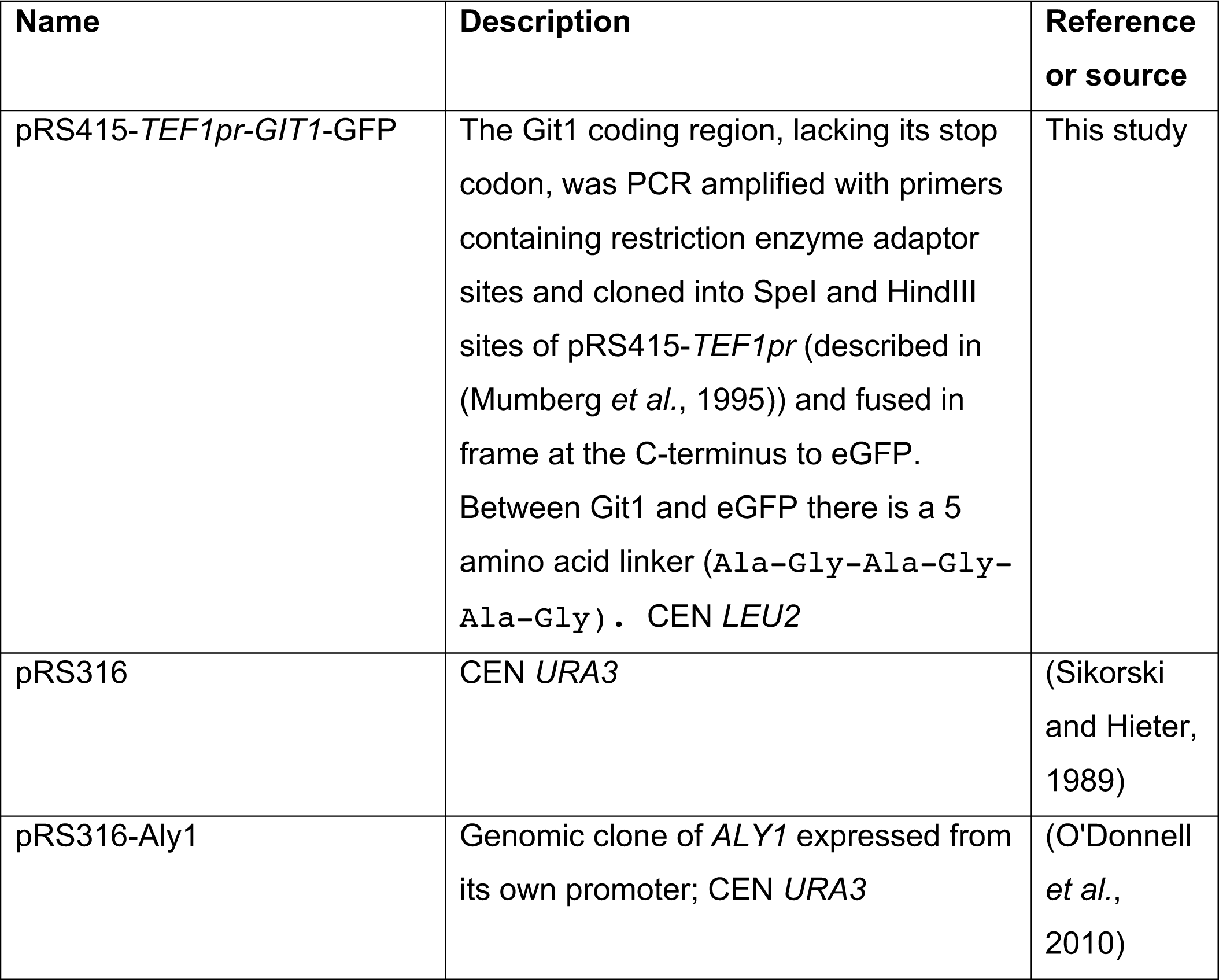
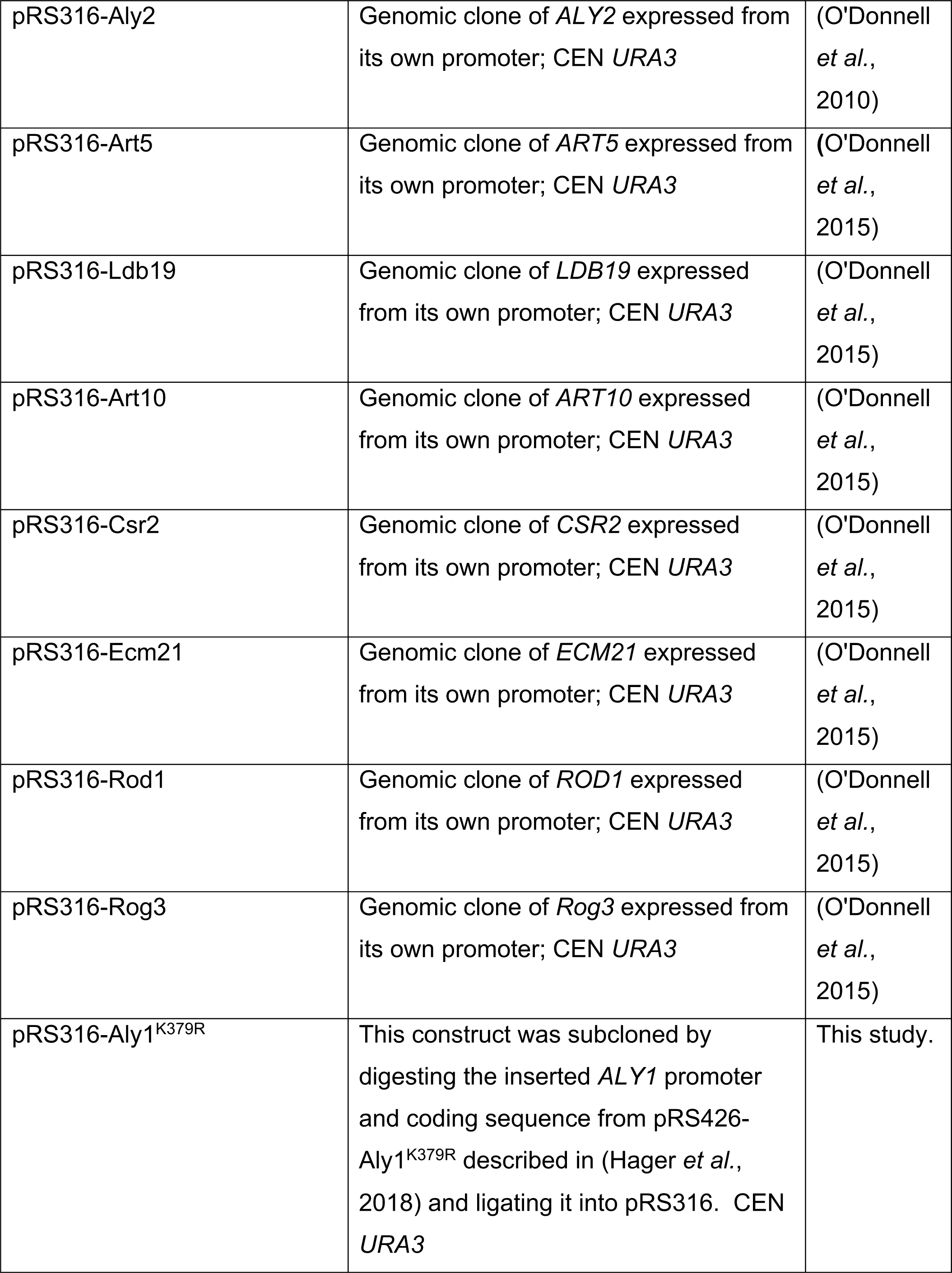
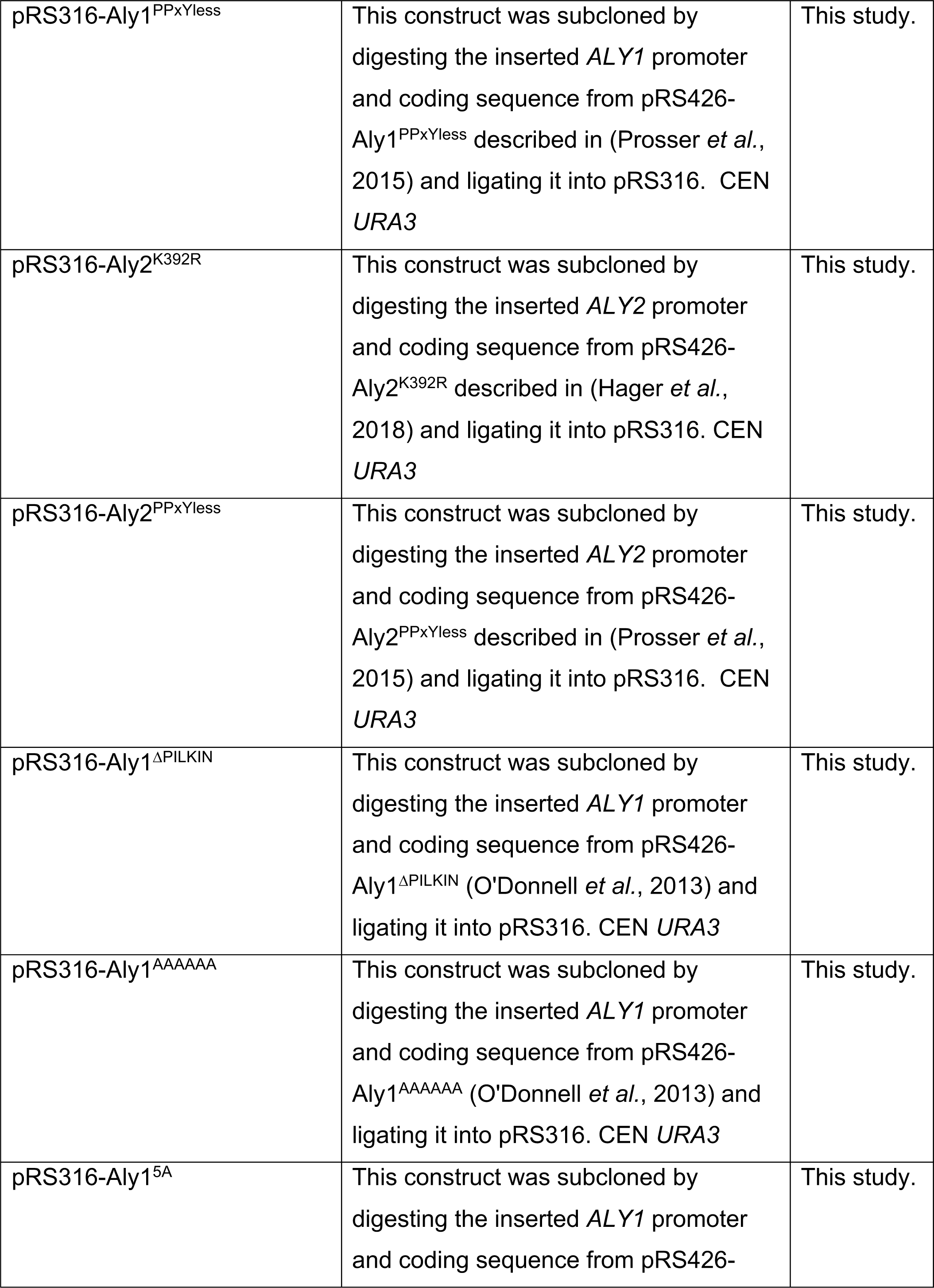
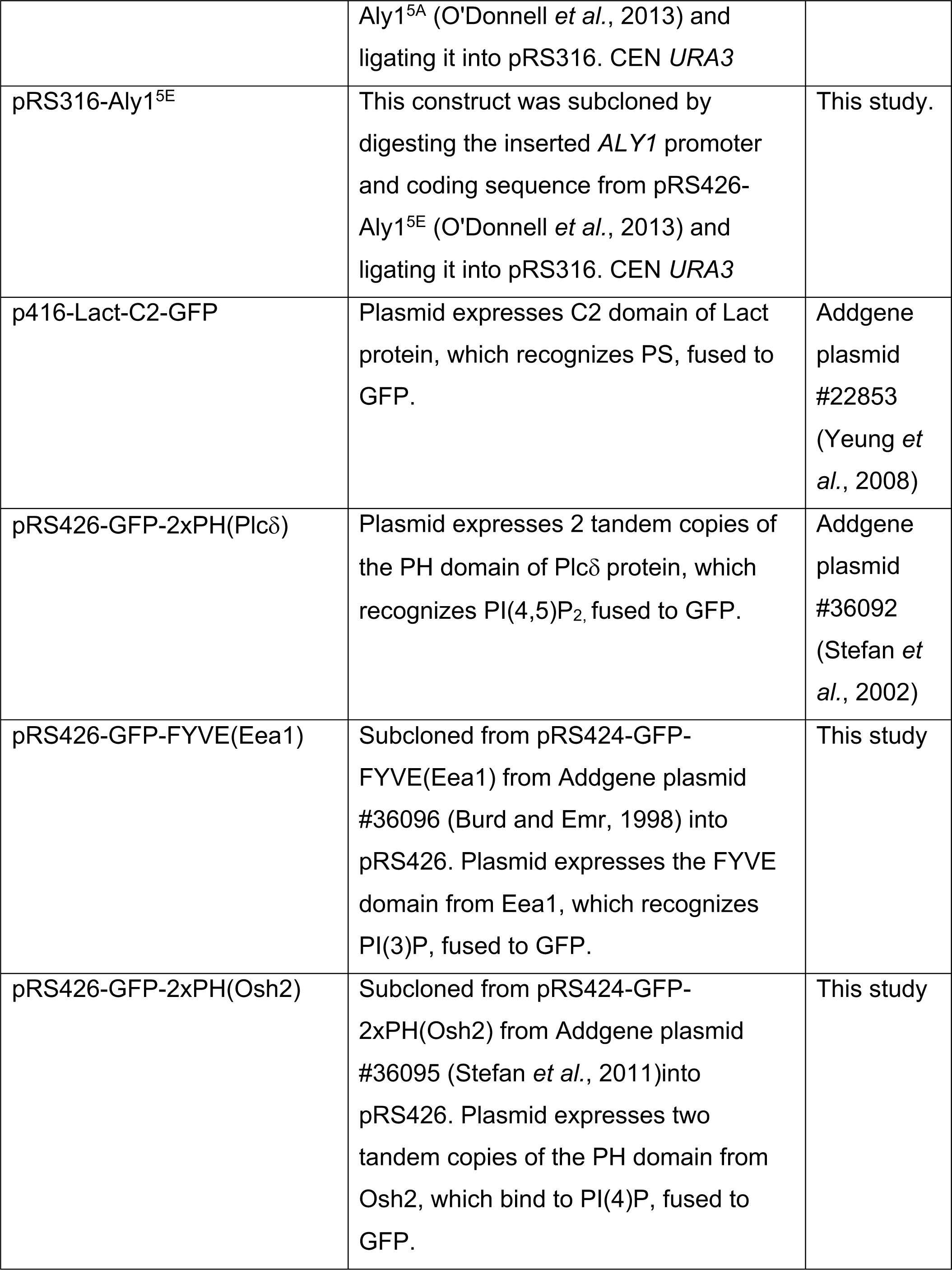
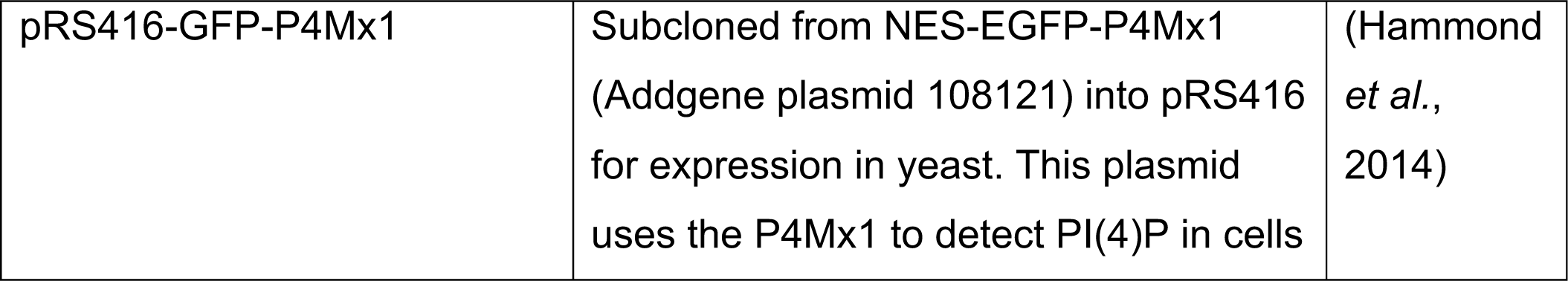
Plasmids.

### Serial dilution growth assays

Yeast cells were grown overnight in liquid culture at 30°C in either SC or YPD medium as appropriate. The saturated cultures absorbance at 600 nm (A_600_) was measured and a starting concentration of A_600_ = 1.0 (which corresponds to approximately 1.0 x 10^7^ cells/ml) was generated. This starting density was diluted in 5-fold series, plated onto the indicated medium and grown for 3-6 days as indicated at 30°C. Plate images were captured on an HP flatbed scanner or using a Chemidoc XRS+ imager (BioRad, Hercules, CA) and all were evenly modified in Photoshop (Adobe Systems Incorporated, San Jose, CA). For GPI containing plates, the GPI was obtained from the Patton-Vogt lab and the final concentration used is indicated in each figure. The myriocin (EMD Millipore, Burlington, MA) and aureobasidin (Takara Contech, Shiga, Japan) concentrations used are indicated in each figure panel.

### Yeast protein extraction and immunoblot analyses

Whole cell extracts of yeast were made using the trichloroacetic acid (TCA) extraction method described in (Hager *et al*., 2018), as modified from (Volland *et al*., 1994). Extracts were resolved by SDS-PAGE and proteins were identified using immunoblotting with mouse anti-GFP antibody (Santa Cruz Biotechnology, Santa Cruz, CA) and a rabbit anti-Zwf1 (glucose-6-phosphate dehydrogenase, referred to as G6PDH) antibody (Sigma, St. Louis, MO) as a protein loading control. Anti-mouse or anti-rabbit secondary antibodies conjugated to IRDye-800 or IRDye-680 (Li-Cor Biosciences, Lincoln, NE) were detected on an Odyssey^TM^ FC infrared imaging systems (Li-Cor Biosciences).

### Fluorescent microscopy

To assess Git1-GFP or GFP-tagged phospholipid probe localization and/or abundance cells were grown overnight to saturation in SC medium, inoculated into fresh SC medium at an A_600_ of 0.2 and grown at 30°C until they reached mid-log phase (A_600_ = 0.5-0.7). Where indicated, cells were treated with 50 μM glycerophosphoinositol (GPI) for the indicated time prior to imaging. Cells treated with latrunculin A (LatA) (Molecular Probes, Eugene, OR) were incubated with 200 μM LatA or an equivalent volume of the dimethyl sulfoxide (DMSO) vehicle control for the indicated time. To visualize vacuoles, cells were stained with 250 μM Cell Tracker Blue CMAC (7-amino-4-chloromethylcoumarin) dye (Life Technologies, Carlsbad, CA). Cells were then plated onto 35 mm glass bottom microwell dishes that were poly-d-lysine coated (MatTek Corporation, Ashland, MA) and imaged using a Nikon Ti inverted microscope (Nikon, Chiyoda, Tokyo, Japan) outfitted with an Orca Flash 4.0 cMOS camera (Hammamatsu, Bridgewater, NJ) and a 100x objective (numerical aperture, 1.49). Image acquisition was controlled using NIS-Elements software (Nikon) and all images within an experiment were captured using identical settings. Images were cropped and adjusted equivalently using Photoshop software (Adobe Systems Incorporated).

### Image quantification and statistical analyses

For all manual fluorescence quantification analyses, fluorescence intensity was measured using Image J software (National Institutes of Health, Bethesda, MD). For cell surface fluorescence quantification, a 2-pixel thick line was hand drawn over the plasma membrane (PM) signal and the mean pixel intensity measured with mean background pixel intensity subtracted (as described in (O’Donnell *et al*., 2015)). For vacuolar fluorescence quantification, the vacuole profile was defined by using images captured of cells stained with CMAC dye (Life Technologies) which was then overlaid on the GFP images and the GFP signal within the vacuole mask was measured using Image J (NIH). The mean background fluorescence intensity was subtracted for the vacuole measures and each vacuole measure was paired to the corresponding plasma membrane quantification. Prism software (GraphPad Software, San Diego, California) was then used to perform Kruskal-Wallis statistical analyses with Dunn’s post hoc test. In all cases, significant p values from these tests are represented as: *, p value <0.1; **, *p* value <0.01; ***, *p* value <0.001; ns, *p* value >0.1 or in some instances where indicated †††, p value <0.001.

For our automated quantification pipeline, we developed an R program called *CellQuant*. The program detects the plasma membrane of cells using a Gaussian blur and simple threshold on the grayscale GFP image. An adaptive threshold based on the minimum plasma membrane radius is then used on the grayscale CMAC vacuole stain to help identify the vacuoles and constrain their size to features that should be smaller than the cell itself. Cells with no detectable vacuolar region are discarded and the fluorescent signal is measured in the GFP channel. The program further pairs the PM fluorescent output measures with the vacuoles and returns the PM/vacuole fluorescent ratios, which are used extensively in this paper. The source code for this pipeline is available at www.odonnelllab.com and https://github.com/sah129/CellQuant. In addition, the Docker image for the program is hosted on Docker Hub at https://hub.docker.com/r/odonnelllab/cellquant, with command-line installation instructions available on GitHub. There are tutorials that can assist individuals who would like to use this pipeline in setting it up on their own systems and we are happy to help users in applying this approach to their own data sets as well.

### Radiolabeled [^3^H]-GPI transport assays and conversion to [^3^H]-PI

For assays monitoring endogenous Git1 activity in the presence or absence of α-arrestins, BY4741 and *aly1*Δ *aly2*Δ cells (biological triplicates of each) were grown aerobically overnight at 30°C in yeast nitrogen base (YNB) medium altered to lack inositol and to contain a low level (200μM) of KH_2_PO_4_ inorganic phosphate and 2% glucose. Amino acid concentrations were as described in (Hanscho *et al*., 2012). These conditions were needed to allow for robust expression of the endogenous Git1 transporter. The next day, single cultures were used to inoculate two fresh cultures containing either 100μM (low) or 10mM (high) inorganic phosphate at an A_600_ of 0.2. All cultures were allowed to reach an A_600_ of 0.5, at which point they were harvested and assayed for transport activity as described below.

Transport assays to monitor endogenous Git1 transport or *TEF1pr*-Git1-GFP transport of GPI were performed as described in (Surlow *et al*., 2014). For the Git1-GFP assays, cells were precultured in high phosphate medium only, as described above, as low phosphate medium was not required to induce expression of Git1. Cells were then harvested by filtration, washed with water and resuspended in 100mM citrate buffer [pH 5] to an A_600_ = 5.0. The assays were initiated by adding 50μl of 75μM [^3^H]-GPI to 200μl of cell suspension. After a 4-minute incubation at 30°C, the assays were stopped by the addition of ice cold distilled, deionized water. Samples were then filtered over Whatman GFC filters (Whatman number 1822-025), washed with 10ml of ice cold distilled, deionized water, and resuspended in 10 ml of scintillation fluid for radioactivity counts. A Student’s t-test was used to assess significance and the data from three biological replicates are shown with each having two technical replicates performed (6 replicates in total). GPI was purchased from Sigma Chemical Company, Tritium labeled GPI ([^3^H]-inositol-GPI) was produced through the deacylation of phosphatidyl[^3^H]-myo-inositol purchased from American Radiolabeled Chemicals, as described (Hama *et al*., 2000).

For assessments of the conversion of GPI to PI after uptake (as in Table 1 and Figure 10C), wild-type or *aly1*Δ *aly2*Δ cultures (biological triplicate) were grown aerobically overnight at 30°C in YNB medium altered to lack inositol and contain a low level of KH_2_PO_4_ (200μM) and 2% glucose. Amino acid concentrations were as described (Hanscho *et al*., 2012). The medium lacked inositol (I-) and contained a low KH_2_PO_4_ level to induce expression of the *GIT1* transporter for GPI uptake. The overnight cultures were diluted to log phase in fresh media containing a high level of KH_2_PO_4_ (10 mM) and allowed to double. To begin labeling, 4 ml cultures normalized to OD_600_ =0.2 were spiked with 5µM [^3^H]-GPI (100,00 cpm/ml). The cultures were harvested following aerobic growth at 30°C for 5 hours. A membrane fraction, a TCA-extractable intracellular fraction and an extracellular fraction were isolated as described previously (Surlow *et al*., 2014). The percentages of radioactive counts in each fraction were determined using a liquid scintillation counter. Lipids were extracted from the membrane fraction and separated by thin layer chromatography as described previously (Surlow *et al*., 2014). Radioactive (American Radiolabeled Chemicals Inc.) and nonradioactive (Avanti Polar Lipid) phosphatidylinositol (PI) standards were used to verify the migration of PI on the TLC plate. Tritium labeled GPI ([^3^H]-inositol-GPI) was produced through the deacylation of phosphatidyl[^3^H]-myo-inositol, as described in (Hama *et al*., 2000).

**FIGURE 10.**
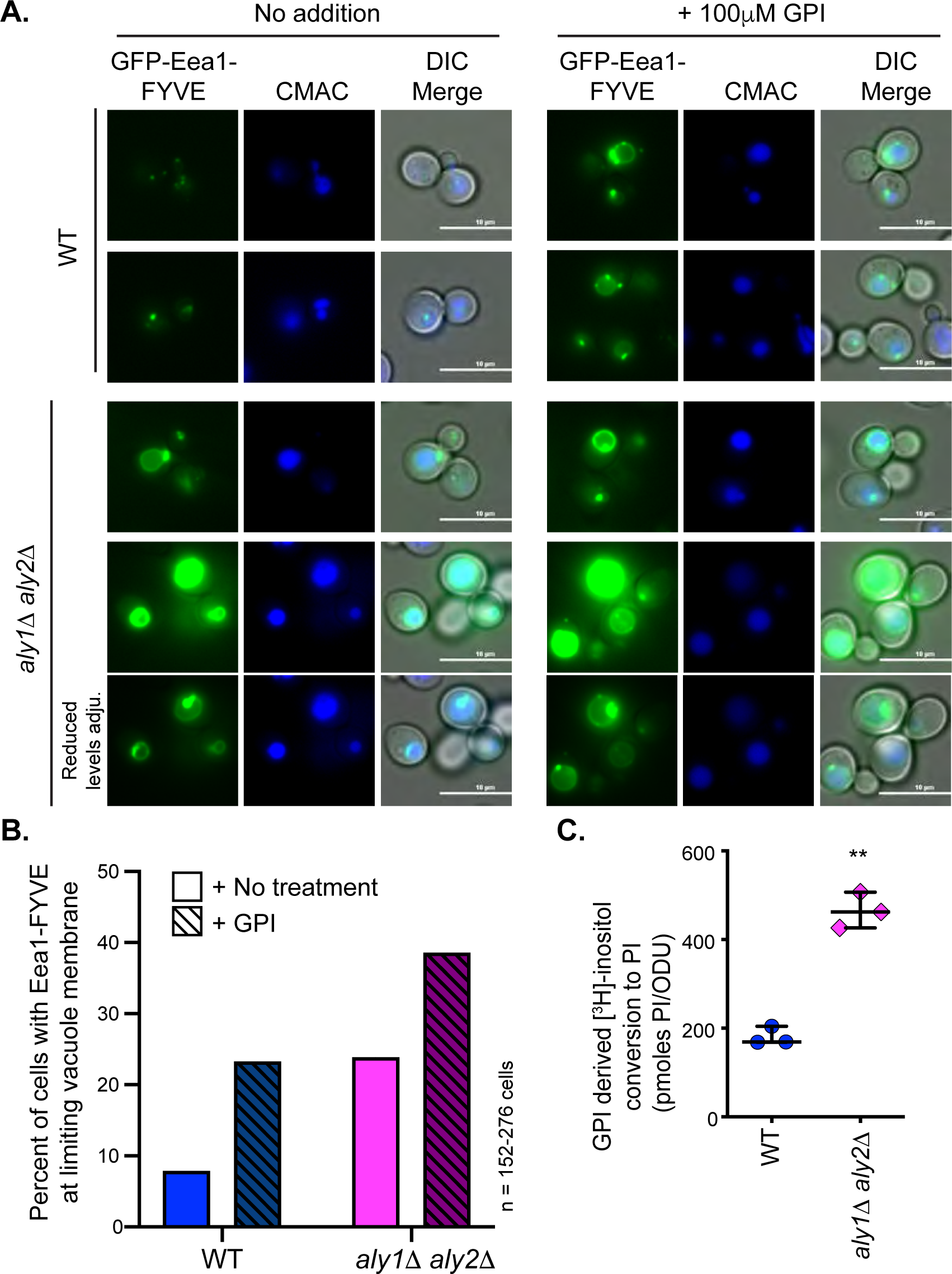
GPI uptake is converted into PI and increases PI3P at the limiting membrane of the vacuole. *A*, The indicated cells were transformed with the plasmid expressing a GFP-fused FYVE domain of Eea1, which marks PI(3)P. Cells were grown to mid-log phase and either untreated (no addition) or treated with 100μM GPI for 2 h, stained with CMAC and imaged by fluorescence microscopy. *B*, Images of cells captured in A were evenly adjusted. Cells with evident GFP staining of vacuolar limiting membrane were counted in comparison to the total number of vacuoles in the image, as assessed using the CMAC stain. The percentage of cells with limiting membrane fluorescence is presented for each treatment. All cells in at least 3 images per condition were counted for an n of 150 at minimum per condition. *C*, Radiolabeled ^3^H-GPI uptake via the endogenous Git1 transporter and conversion to ^3^H-PI was measured in wild-type or *aly1*Δ *aly2*Δ cells. Cells were grown under low phosphate conditions to induce *GIT1* expression in the presence of 5 μM [^3^H-inositol]-GPI, and intracellular and membrane fractions were isolated. The conversion of ^3^H-GPI into ^3^H-PI was monitored by using TLC and liquid scintillation counting. The graph presented represents three independent biological replicates that were measured and a Student’s t-test was used to assess significance. A more detailed breakdown of these results is presented in Table 1 where both the water-soluble and membrane fractions are compared.

**FIGURE 11:**
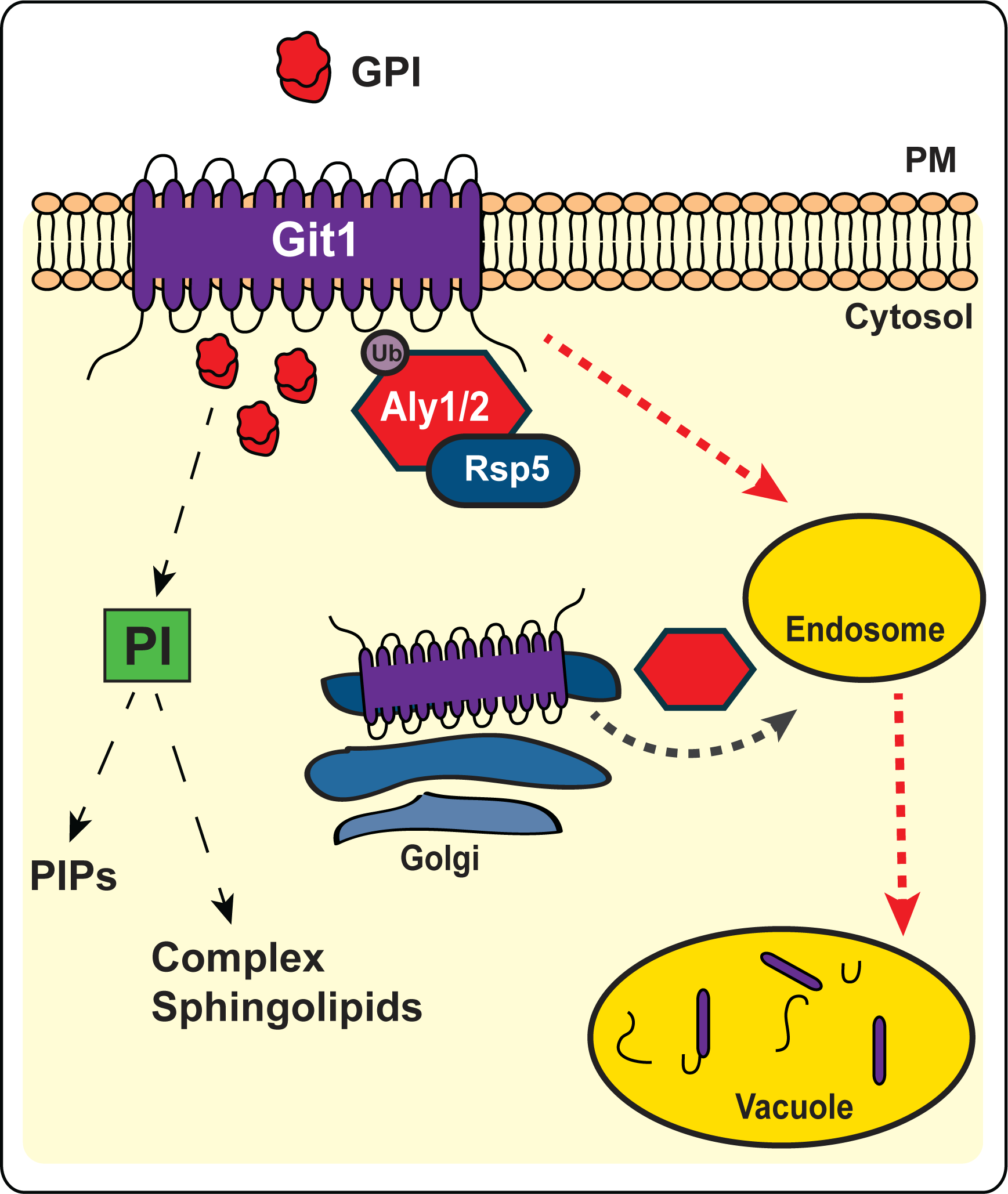
Model of α-arrestin regulation of Git1 trafficking. α-Arrestins are responsible for trafficking of Git1 to the vacuole during both steady-state conditions, where Aly2 is predominantly active, and in response to its ligand GPI, where Aly1 seems to play a more important role. There is also an intracellular sorting pathway that is independent of endocytosis that contributes to Git1 trafficking to the vacuole and this pathway is also α-arrestin dependent (indicated by grey dashed arrow). Association with the ubiquitin ligase Rsp5 is required for α-arrestin-mediated trafficking of Git1. The endocytic pathway (indicated by red dashed lines) appears to be clathrin-dependent as disruption of clathrin-associated factors, like End3 and AP-2, leads to retention of Git1 at the cell surface. Uptake of GPI via Git1 may alter the sphingolipid balance in cells and seems to directly influence the distribution of membrane phospholipids, including elevating PI3P on the vacuolar membrane.

## Abbreviations

a.u.: arbitrary units
CDP: cytidine diphosphate
DMSO: Dimethyl sulfoxide
GPI: glycerophosphoinositol
LatA: Latrunculin A
PI: phospatidylinositol
PI(3)P: phosphatidylinositol-3-phosphate
PI(4)P: phosphatidylinositol-4-phosphate
PI(3,5)P_2_: phosphatidylinositol-3,5-bisphosphate
PI(4,5)P_2_: phosphatidylinositol-4,5-bisphosphate
PS: phosphatidylserine

---

**SUPPLEMENTAL FIGURE 1. Impact of the Buls on GPI sensitivity and Git1 trafficking.** *A*, Growth of serial dilutions of the cells indicated containing a *LEU2*-marked *TEF1pr*-*GIT1*-GFP plasmid on SC medium lacking leucine and containing 25μM or 50μM GPI. Growth is shown at 2 days after incubation at 30°C. *B*, Fluorescence microscopy of the indicated cells containing Git1-GFP expressed from the *TEF1* promoter. *C*, The PM and vacuolar fluorescence intensities from the cells depicted in B were quantified using our automated quantification pipeline and the distributions of PM/intracellular fluorescence ratios in arbitrary units (a.u.) were plotted as scatter plots. The horizontal midline in black represents the median and the 95% confidence interval is represented by the white error bars. Kruskal-Wallis statistical analyses with Dunn’s post hoc test were performed and statistical significance compared to the wild-type cells is indicated with asterisks. A yellow dashed line is provided that represents the median of wild-type cells to facilitate comparisons.

**SUPPLEMENTAL FIGURE 2. Assessing function of the Git1-GFP construct and regulation of endogenous Git1 by Aly1 and Aly2.** *A*, Growth of serial dilutions of the cells indicated either containing a *LEU2*-marked vector control (vec.) or expressing a *LEU2*-marked *TEF1pr*-*GIT1*-GFP plasmid on medium lacking leucine, glucose, and inositol and containing either high phosphate (10 mM), low phosphate (200μM) or no phosphate and supplemented with 100μM GPI where indicated. Growth is shown after incubation at 30°C. *B-C*, Automated quantification of the imaging data shown in Figure 3C and 3D, respectively. These data, while following the same trend as the manual quantification of these data presented in Figures 3E and 3F, respectively, demonstrate how the automated pipeline can underestimate the number of cells with no fluorescent signal at the plasma membrane, as is the case for these wild-type cells.

**SUPPLEMENTAL FIGURE 3. α-Arrestin-mediated trafficking of Git1 in LatA and GPI treated cells.** *A*, As done in Figure 5D-E, the cells indicated containing Git1-GFP expressed from the *TEF1* promoter were cultured in SC media lacking leucine and containing 200μM latrunculin A or an equal volume of DMSO vehicle for 120 min prior either addition of 50μM GPI or no treatment, and incubated a 120 min more before being stained with CMAC and imaged by fluorescence microscopy. These images were quantified using our automated quantification pipeline only (and this is the first time the manual quantification of these data is not also performed) and the distributions of PM/vacuolar fluorescence ratios in arbitrary units (a.u.) were plotted as scatter plots. The horizontal midline in black represents the median and the 95% confidence interval is represented by the white error bars. A blue dashed line represents the median ratio for untreated wild-type cells and a pink dashed line represents the median ratio for untreated *aly1*Δ *aly2*Δ cells to facilitate comparisons. Kruskal-Wallis statistical analyses with Dunn’s post hoc test were performed within each graphing group and statistically significant differences are marked with asterisks or indicated by ns if not significant.

**SUPPLEMENTAL FIGURE 4. The impact of CN-regulation mutants in Aly1 on Git1 steady-state trafficking.** *A*, WT or *aly1Δ aly2Δ* cells containing Git1-GFP expressed from the *TEF1* promoter and either a pRS316 empty vector expressing nothing or plasmid expressing the indicated α-arrestins were stained with CMAC and imaged by fluorescence microscopy. *B*, The PM and vacuolar fluorescence intensities from the cells depicted in A were quantified using our automated quantification pipeline and the distributions of PM/intracellular fluorescence ratios in arbitrary units (a.u.) were plotted as scatter plots. The horizontal midline in black represents the median and the 95% confidence interval is represented by the white error bars. Kruskal-Wallis statistical analyses with Dunn’s post hoc test were performed and statistical significance compared to the *aly1*Δ *aly2*Δ cells with vector is indicated with asterisks or as not significant (ns) where applicable. In each case, a yellow dashed line is provided that represents the median of the *aly1*Δ *aly2*Δ cells containing a vector control to facilitate comparisons.

**SUPPLEMENTAL FIGURE 5. Altered trafficking changes sphingolipid balance.** *A*, Growth of serial dilutions of the cells indicated on YPD medium containing the sphingolipid regulatory drugs indicated is shown at 2 days after incubation at 30°C.

**SUPPLEMENTAL FIGURE 6.** *A*, The indicated cells were transformed with the plasmid expressing GFP-fused Plc*δ*, which marks PI(4,5)P_2_. Cells were stained with CMAC and imaged by fluorescence microscopy. *B*, The GFP fluorescence at the plasma membrane from the images in A was quantified using our automated image analyses pipeline. The quantified fluorescence intensities (arbitrary units) are presented as scatter plots. The horizontal midline in black represents the median and the 95% confidence interval is represented by the black error bars. Kruskal-Wallis statistical analyses with Dunn’s post hoc test were performed and statistical significance compared to the wild-type cells is indicated as not significant (ns) in all cases. A yellow dashed line is provided that represents the median of wild-type cells to facilitate comparisons. *C*, The indicated cells were transformed with the plasmid expressing a GFP-fused P4Mx1, which marks PI(4)P. Cells were stained with CMAC and imaged by fluorescence microscopy. It should be noted that similar results were obtained using the pRS426-GFP-2xPH domain of Osh2 to assess PI(4)P levels in these same strains (not shown).

## Notes

### Competing Interest Statement

The authors have declared no competing interest.

