## Supplementary material for "Alpha-arrestins Aly1 and Aly2 regulate trafficking of the glycerophosphoinositol transporter Git1 and impact phospholipid homeostasis": Robinson et al Supplemental

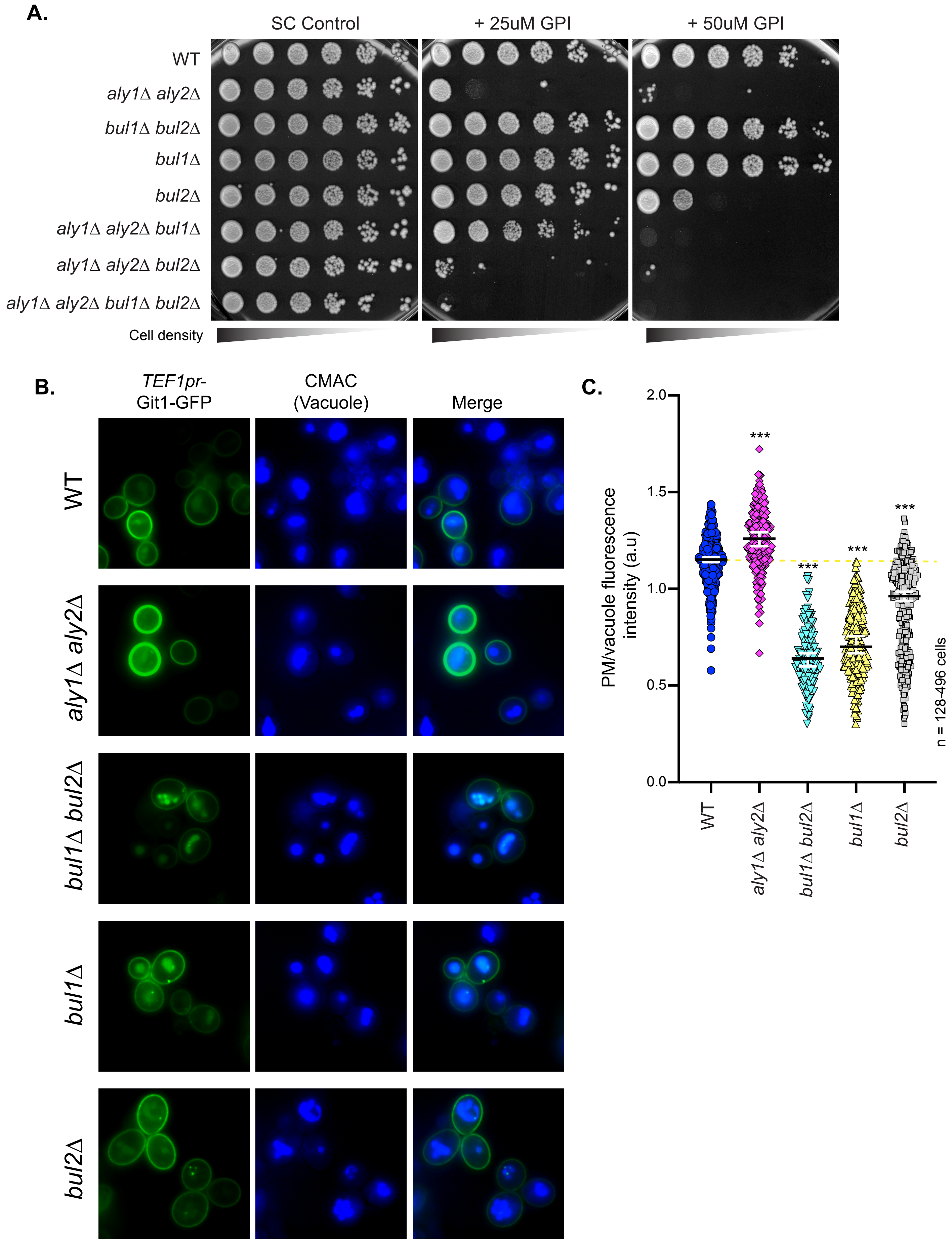


**SUPPLEMENTAL FIGURE 1**

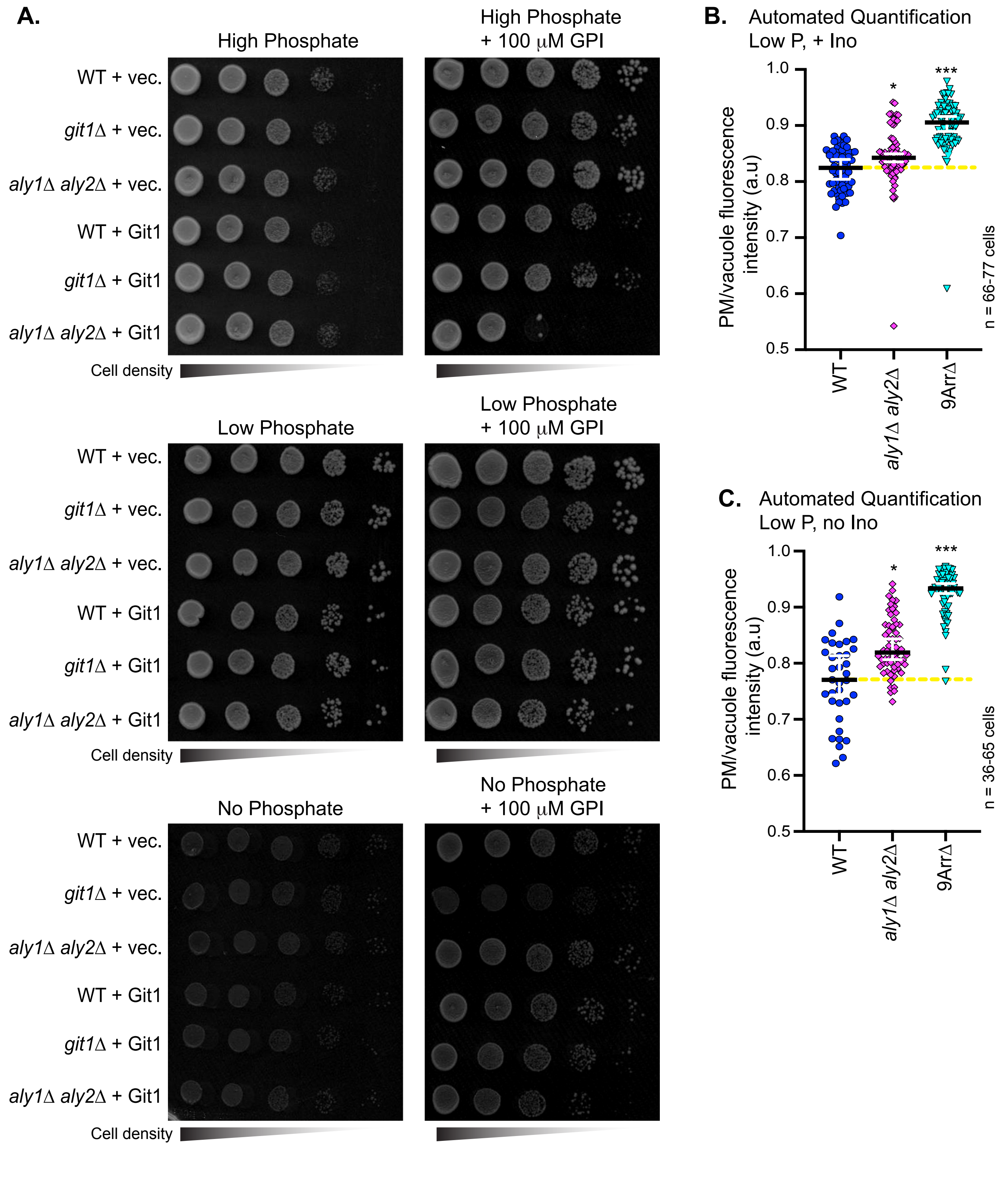


**SUPPLEMENTAL FIGURE 2**

**A.**

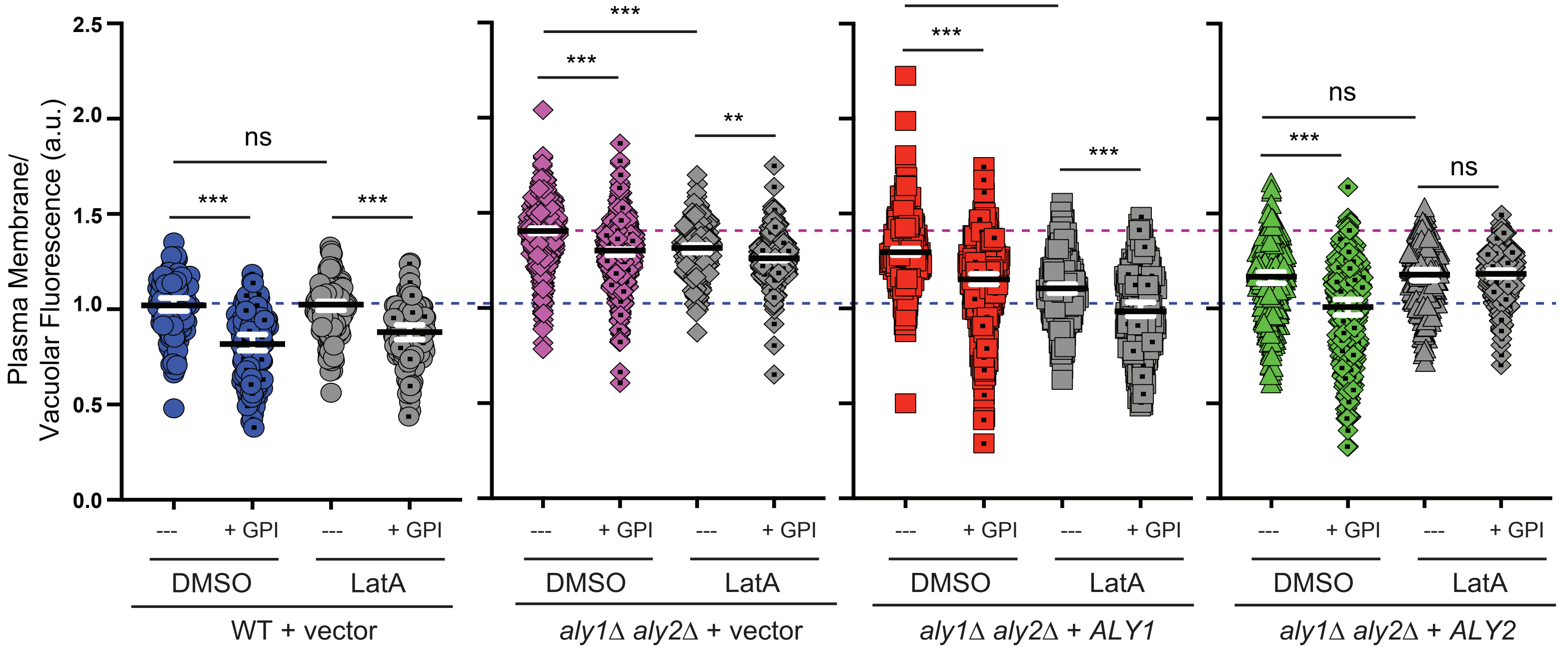


n = 124-520 cells per condition

**SUPPLEMENTAL FIGURE 3**

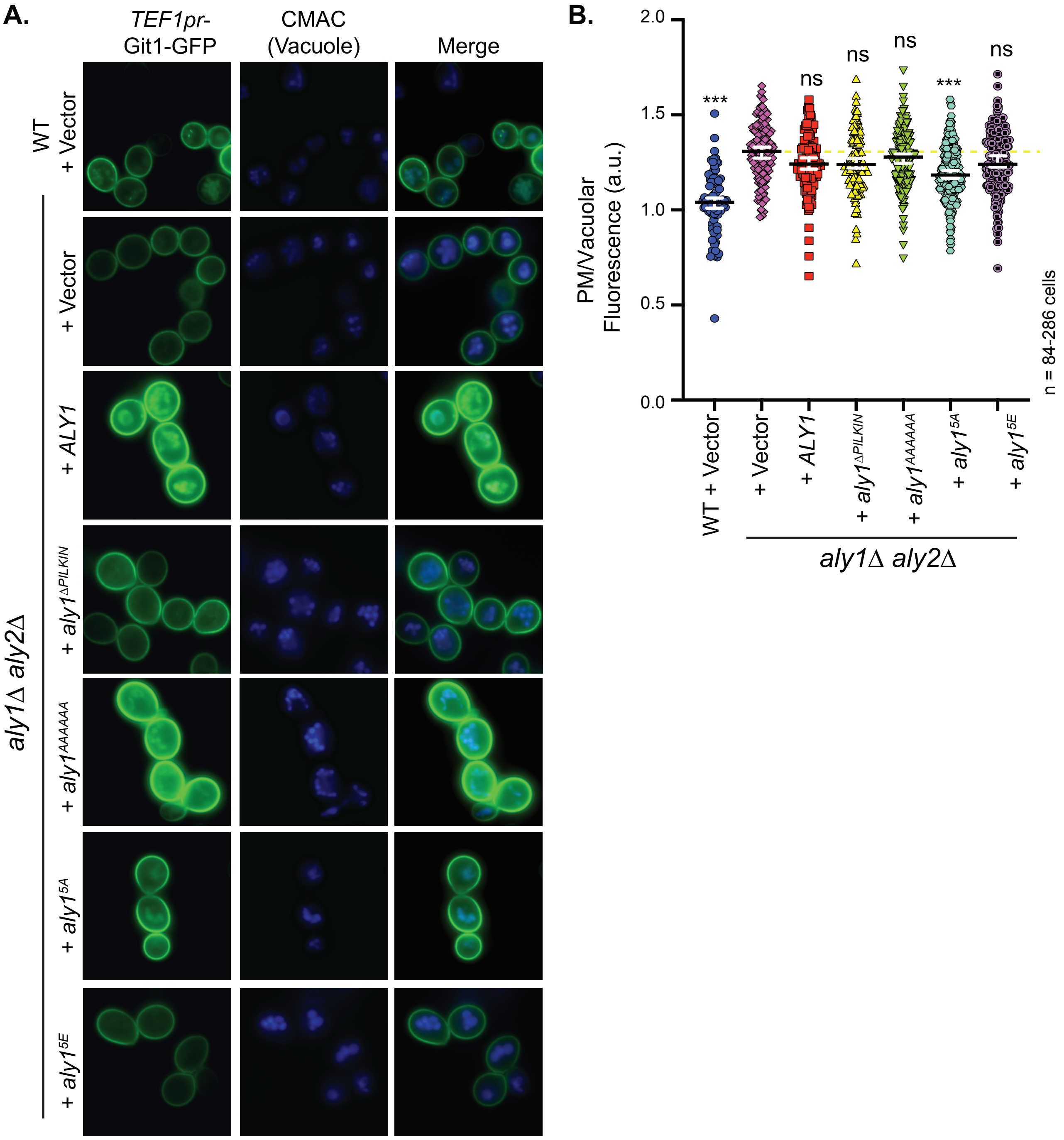


**SUPPLEMENTAL FIGURE 4**

**A.**

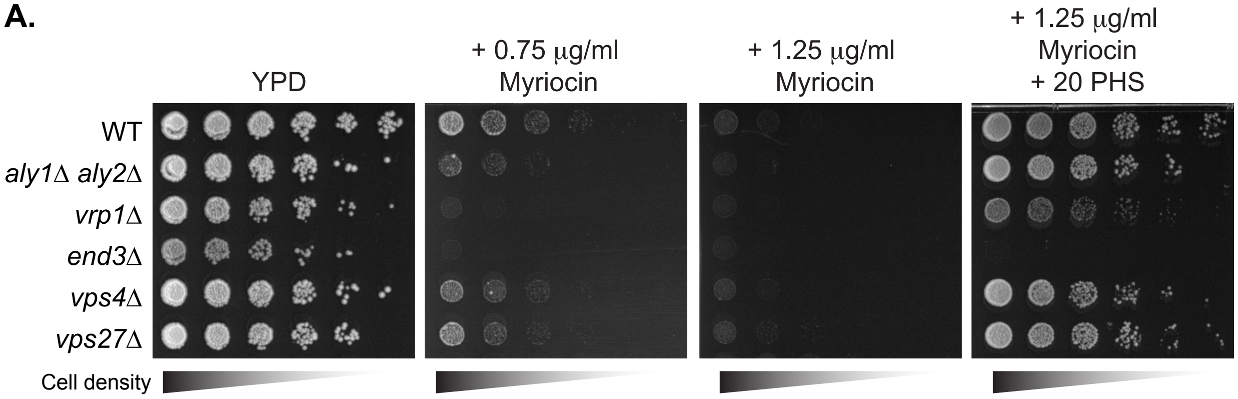


**SUPPLEMENTAL FIGURE 5**

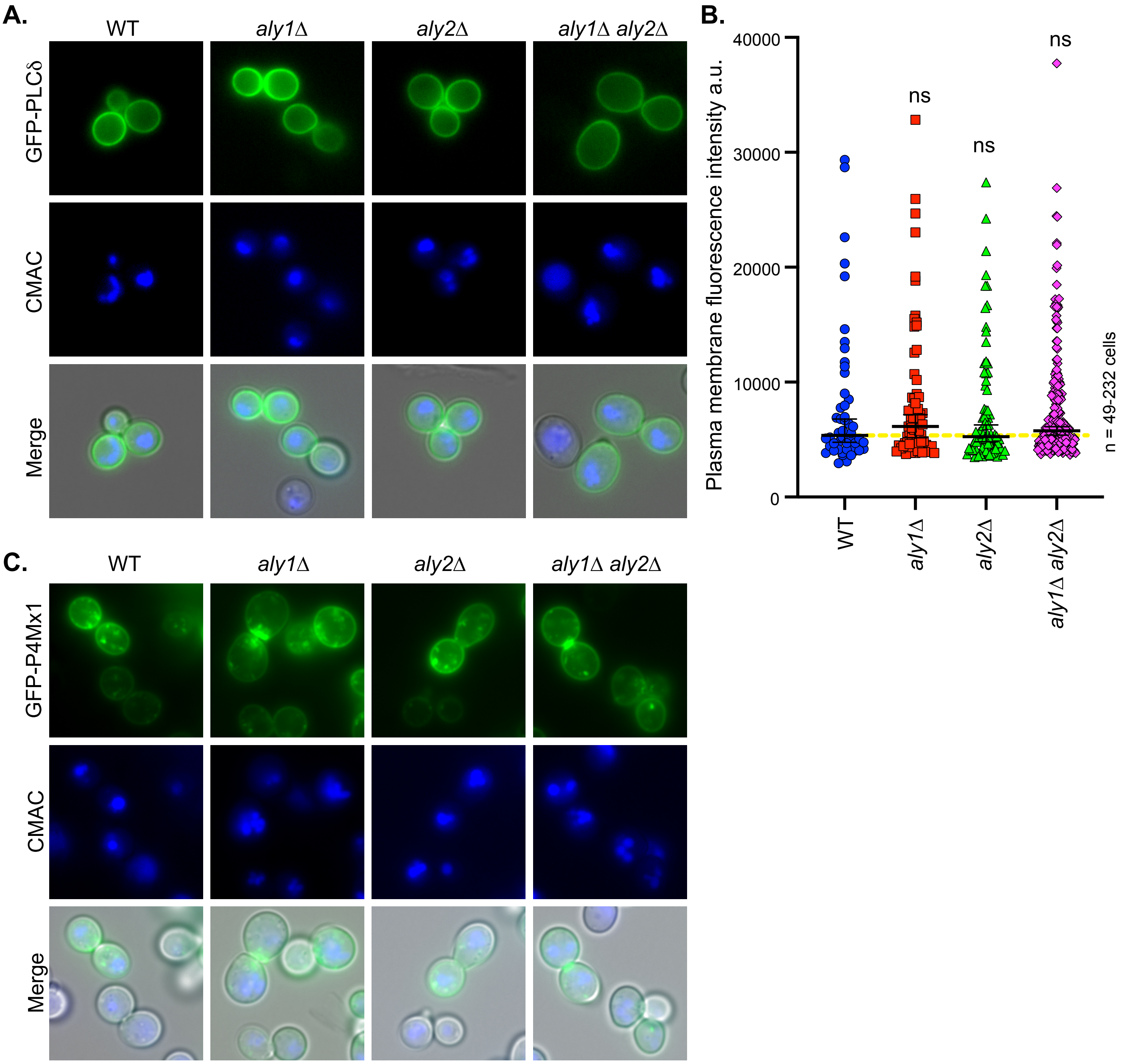


**SUPPLEMENTAL FIGURE 6**
